# Data Processing Pipeline for Multiband Diffusion-, T1- and Susceptibility Weighted MRI to Establish Structural Connectivity of the Human Basal Ganglia and Thalamus

**DOI:** 10.1101/2020.03.06.981142

**Authors:** Irtiza A. Gilani, Kader K. Oguz, Huseyin Boyaci, Katja Doerschner

## Abstract

The basal ganglia and thalamus play an important role in cognition, procedural learning, eye movements, control of voluntary motor movements, emotional control, habit development, and are structures that are severely impacted by neurological disorders such as Parkinson’s disease, or Tourette syndrome. To understand the structural connectivity of cortical and subcortical circuits in the healthy human brain could thus be of pivotal importance for detecting changes in this circuitry and to start early intervention, to assess the progress of movement rehabilitation, or the effectiveness of therapeutic approaches in neuropsychiatry. While conventional clinical magnetic resonance imaging (MRI) is able to provide detailed information about connectivity at the macro level, the sensitivity and specificity of such structural imaging methods put limits on the amount of detail one can obtain when measuring *in vivo* connectivity of human basal ganglia and thalamus at routine clinical magnetic field strengths. In contrast, the multiband diffusion echo planar imaging method, which acquires multiple slices simultaneously, enables high resolution imaging of these abovementioned brain structures with only short acquisition times at 3-Tesla and higher magnetic field strengths. To unleash the greater potential of information embedded in data acquired with this technique, complementary data processing pipelines are required. Here, we use a protocol composed of multiband diffusion-, T1- and susceptibility weighted data acquisition sequences and introduce an associated pipeline based on combined manual and automated processing. The design of this data processing pipeline allows us to generate comprehensive *in vivo* participant-specific probabilistic patterns and visualizations of the structural connections that exist within basal ganglia and thalamic nuclei. Moreover, we are able to map specific parcellations of these nuclei into sub-territories based on their connectivity with primary motor-, and somatosensory cortex. This data processing strategy enables detailed subcortical structural connectivity mapping which could benefit early intervention and therapy methods for human movement rehabilitation and for treating neuropsychiatric disorders.

## Introduction

The basal ganglia are part of several neuronal pathways that control emotional, motivational, and cognitive functions [Weyhenmer et al., 2007; Niv et al., 2007; Stocco et al., 2010]. They play a major role in the functioning of the complex extrapyramidal motor system, and functional abnormalities of the basal ganglia are associated with many movement disorders, such as Parkinson’s and Huntington’s disease, abulia or dystonia [Marin and Wilkosz, 2005; Baker et al., 2013; Cameron et al., 2010; Crossman, 2000].

The basal ganglia create a complex network with several cortical and subcortical areas [Takada et al., 1998; Bar-Gad and Bergman, 2001]. Graybiel et al. [2000] suggest that basal ganglia structures operate as part of the recurrent circuits (loops) with the cerebral cortex and form several cortico-basal ganglia loops. Previous models of cortico-basal ganglia-thalamic-cortex connectivity suggest that parallel pathways play a role in regulating sensorimotor, associative, and limbic information processing [Haber, 2003; Haber and Calzavara, 2009; Cohen and Frank, 2009], however, McFarland and Haber [2002] proposed that such parallel connectivity patterns are not one-way loops and the pathway back to cortex has one component that reinforces each cortico-basal ganglia circuit and one component that relays information between circuits.. In general, these basal ganglia connectivity loops are thought to include connections not only with cortex but also within-, and between basal ganglia and other subcortical structures, such as thalamus and superior colliculus (McHaffie et al., 2005). In addition to reciprocal connections in the cortico-basal ganglia-thalamic circuits that connect regions associated with similar cognitive functions (maintaining parallel networks), there are also non-reciprocal connections (integrative networks) linking regions that are associated with different cortical-basal ganglia-thalamic circuits [Haber and Calzavara, 2009]. Integration of inputs to emotional, cognitive and motor functions may occur via these non-reciprocal connections, for example, between striatum and substantia nigra, between cortico-striatal projections from different functional regions, and between thalamus and subcortical regions via different thalamo-subcortical projections.

Given the complexity of basal ganglia network, there is need to build appropriate data acquisition methods and processing tools capable to measure and visualize the topology of this network in order to gain a deeper understanding of these circuits. Previous research in this area has been mainly based on animal data [Kelly and Strick 2004; Francois et al., 2004; Joel and Weiner 1997; Kuo and Carpenter 1973; Rouiller et al., 1994], patient data [Joel 2001; Mallet et al. 2007], averaged human brain data [Zhang D et al., 2010; Klein JC et al., 2010; Lehericy S et al., 2004; Menke RA et al., 2010; Aravamuthan BR et al., 2007; Draganski B et al., 2008], post-mortem histological data [Gallay et al., 2008; Dyrby et al., 2007], and translational connectivity analysis from non-human primates to humans [Mars et al., 2011]. Although these studies have made major contributions, there is still a lack of consensus on data processing strategies which use ultra-large scale multi-contrast human brain data sets acquired with state-of-the-art imaging methods and provide comprehensive description of *in vivo* human subcortical circuitry. Here we provide proof-of-concept of pipeline design by focusing on identification of detailed connectivity patterns of the human subcortical brain regions using diffusion MRI data acquired with short repetition time (TR) and echo time (TE) at 3T.

Diffusion weighted imaging (DWI) is a structural magnetic resonance imaging (MRI) technique that enables white matter integrity analysis *in vivo* [Pierpaoli and Basser, 1996], and by combining it with probabilistic tractography the trajectory and microstructural integrity of white matter pathways can be estimated. Researchers have used this approach [e.g. Behrens et al., 2003; Johansen-Berg et al., 2005; Leh et al., 2007] to perform thalamic-, and striatal connectivity-based parcellation to segment brain regions that were not directly detectable by conventional MRI techniques [Klein et al., 2007]. This connectivity-based segmentation of brain structures has also been used for parcellating the basal ganglia and thalamus [Draganski et al., 2008], however, their automated region of interest segmentation did not aim to capture some of the connections, e.g. those to hippocampus, fusiform gyrus or substantia nigra. In our work, we prefer manual segmentation to automated segmentation of regions of interests in order to capture such regions accurately and precisely, in combination with high-resolution and improved diffusion contrast to perform automated connection-based parcellation of basal ganglia and thalamus. Specifically, we employ a rapid DWI acquisition method, with short TR and TE, because this MRI sequence is beneficial for reduction in SNR loss due to T_2_ decay inherent to the diffusion encoding and acquisition processes in MRI [Ugurbil et al. 2013, Sotiropoulos et al. 2013]. Thus, this approach generates minimal SNR per unit time penalty for the data acquired at 3T. Our pipeline is built for utilizing the acquired data’s superior capability of capturing structural connectivity information in the human brain.

Previously, human basal ganglia and thalamus have been manually segmented and parcellated at 7T using the probabilistic tractography approach for processing data (Langelt et al. [2012]). This data processing pipeline is similar to our proposed design, however, our methodology is more focused on sub-region identification at 3T and is not based on T2-weighted MRI data. The authors of of that study employed a high-angular resolution diffusion MRI sequence with an in-plane acquisition acceleration factor=3 and a head gradient inset capable of 80mT/m in 135 msec. In general, the intrinsic SNR of MR images is elevated at ultra-high magnetic fields [Vaughan et al. 2001]. For DWI, however there could be an SNR advantage at 7T (over 3T) only if TE could be kept under 100 ms [Ugurbil et al. 2013]. Therefore, improved DWI signal could only be obtained if the 7T scanner is equipped with 150 to 300 mT/m maximum gradient strength, which in turn would bring TE values well under 50 ms [Ugurbil et al. 2013]. Consequently, Langelt et al. [2012] did not benefit fully from the potential SNR advantages of the 7T scanner.

In this study, we used a 3T scanner without expensive gradients and ultra-high field (>3T) technology. Our approach may thus be more suitable for conditions common in the clinical setting. We acquired human brain data using a simultaneous multi-slice acquisition based DWI sequence [Setsompop et al., 2012]. A reduction in TR was achieved by simultaneous excitation of 3 slices, which results in a 3-fold reduced TR (compared with standard DWI sequences) and a less than 100 ms TE. The proof-of-concept of a semi-automated probabilistic tractography-based data processing pipeline is provided by generating subject-specific comprehensive structural connectivity pathways of human basal ganglia and thalamus. Taken together, we demonstrated that this pipeline can be integrated with diffusion MRI data acquired with short TR and TE at 3T yielding a highly detailed connectivity map and parcellation of human subcortical brain regions.

## Methods

### Participants

Five healthy subjects (4 females and 1 male) provided informed written consent prior to participating in this study. All subjects (27.6**±**7.43 years) were right handed, and none of them had a history of brain abnormalities and neurological disorders. The study adhered to the principles put forward by the declaration of Helsinki and was approved by the ethical review committee of Bilkent University, Ankara, Turkey.

### Data acquisition

All subjects were scanned at the National Magnetic Resonance Research Center (UMRAM) in Bilkent University, using a 3T MRI system (Siemens Magnetom Trio, Germany). A 32-channel head coil was used for data acquisition. The MRI protocol was composed of a T1-weighted sequence, a high-resolution susceptibility-weighed imaging (SWI) sequence and a high-resolution DWI sequence, described next.

#### Structural MRI

T1-weighted images were acquired with a Siemens standard 3D magnetization-prepared rapid acquisition gradient-echo (MP-RAGE) sequence. MP-RAGE incorporates an inversion pulse to enhance the T1 weighting and obtains excellent contrast between gray matter (GM) and white matter (WM) in the cortical and subcortical regions of the brain. In this work, MP-RAGE sequence was used with the following parameters: 176 slices; field of view (FoV), 256 ⨯ 224 mm; matrix, 256 ⨯ 256; 1 mm^3^ isotropic voxels; sagittal, phase encoding in anterior/posterior; repetition time (TR), 2600 ms; echo time (TE), 3.02 ms; inversion time (TI), 900 ms; flip angle, 8 degrees; bandwidth, 130 Hz/pixel; echo spacing, 8.9 ms; parallel imaging, GRAPPA with an acceleration factor of 2 along the phase-encoding direction; phase partial Fourier, 6/8; slice partial Fourier, 7/8. The total acquisition time was approximately 5 min.

Susceptibility-weighed MR imaging (SWI) is a powerful tool to visualize the iron content in the subcortical region of brain, i.e. substantia nigra (SN). SWI data were acquired with a Siemens’ high-resolution axial 3D SWI sequence using the following parameters: FOV, 180 ⨯ 180 ⨯ 224 mm^3^; matrix, 448 ⨯ 448; 72 slices (20% slice gap); resolution, 0.4 ⨯ 0.4 ⨯ 2 mm^3^; axial, phase encoding in anterior/posterior; TR, 28 ms; TE, 20 ms; flip angle, 15 degrees; bandwidth, 120 Hz/pixel; parallel imaging, GRAPPA with an acceleration factor of 2 along the phase-encoding direction. It is important to note that only the subcortical region was covered during the SWI acquisition, not the whole brain. The total acquisition time was approximately 5 min.

#### Diffusion MRI

Whole brain diffusion-weighted MRI data were acquired with 1.8 ⨯ 1.8 ⨯ 1.8 mm^3^ resolution using 2D multiband multislice spin-echo EPI sequence [Setsompop et al., 2012] and the following parameters: 81 slices; FoV, 192 ⨯ 192 ⨯ 192 mm^3^; matrix size, 108 x 108; TR, 3875 ms; TE, 87.4 ms; flip angles, 78 and 160 degrees; bandwidth, 1494 Hz/pixel; echo spacing, 0.77 ms; excitation pulses durations, 3200 and 7040 µs; phase partial Fourier, 6/8; fat suppression, weak fat saturation; multiband factor, 3; b-value, 1000 s/mm^2^); diffusion gradients were applied along 64 uniformly distributed directions. This leads to a total acquisition time of 4 min 51 sec, which is much shorter than typical diffusion MRI sequences having a TR in the range of 7000-8000 ms.

### Description of Data Processing Pipeline

#### Pre-processing of diffusion MRI data

Pre-processing of diffusion-weighted MR images was performed using the FMRIB’s software library (FSL), version 5.0 (FMRIB, Oxford, UK). The following steps were repeated for each participant’s data. Initially, diffusion-weighted images were corrected for the eddy current distortions using FMRIB’s diffusion tool (FDT), version 3.0. The diffusion gradients were reoriented using the modified values that were generated after the eddy current correction for each diffusion-weighted image. The non-diffusion-weighted image (D0) was extracted from the acquired diffusion data. FSL’s brain extraction tool (BET) version 2.1 was used to extract the brain region without skull from the non-diffusion-weighted and diffusion-weighted images and to obtain the corresponding binary masks. Thereafter, FDT’s BEDPOSTX utility was employed with the option of two crossing fibers enabled for estimating fiber orientations at each voxel.

#### Segmentation of regions-of-interest and transformations

Sixteen subcortical regions-of-interest (ROIs) were manually segmented using structural images (T1-weighted and SWI) of each participant. We used a neuroanatomical atlas [Atlas for stereotaxy of human brain, Schaltenbrandt and Wahren, Nov 1977, Thieme, 2nd edition,ISBN: 9783133937023, 84 pages] to identify and create masks for eight ROIs per hemisphere: caudate nucleus (CN), putamen (Pu), globus pallidus internal (GPi), globus pallidus external (GPe), substantia nigra pars reticulata (SNr), substantia nigra pars compacta (SNc), subthalamic nucleus (STN) and thalamus (Tha).

In order to recheck the identified regions in the T1-weighted images, the CN was traced in the coronal plane by considering the white matter of the anterior limb of the internal capsule as the lateral boundary and the lateral ventricle as the medial boundary. The Pu, lateral to the CN, was also traced in the coronal plane. The internal capsule was considered as the medial boundary and the external capsule was taken into account as the lateral boundary. The corona radiata was marked as the superior boundary and the white matter of the retrolenticular limb of the internal capsule was used as the posterior boundary for tracing Pu. The GP was also marked in the coronal plane using as anterior border where it appeared medially to the Pu. The internal capsule was used to identify the superior and medial borders. The lateral border coexisted with the Pu and the inferior border was defined using the substantia innominata and the anterior commissure. The end of GP slice was marked where the GP vanished medially to the Pu. The region between the GPe and GPi was identified by using the medial medullary lamina as a separating structure.

Additionally, we also compared the basal ganglia areas traced in the coronal plane with the corresponding outlined areas in the transverse and sagittal planes. For that we used axial SWI images to segment SNc and SNr. The SNc was distinguishable from the surrounding tissue, especially the red nuclei, due to its dark grey color in contrast to the adjacent regions, and the region between the red nuclei and SNc was marked as SNr. Coronal SWI images were used to re-check the segmentation of SNr and SNc, since images in this plane allow the evaluation of the separation of the SN region along the lateral-medial axis.

In subsequent steps, all ROI masks were transformed from their native structural space to their corresponding diffusion space. To do this we first aligned T1-weighted and SWI data with non-diffusion-weighted data using FMRIB’s linear registration tool version 6.0, and then used the resulting transformations to warp the ROI masks to the diffusion space (Figure 1). All masks were thresholded with value 0.5. The segmentation of GPe and GPi is shown in Figure 2 A, and the segmentation of the SNc and SNr regions using the SWI data is illustrated in Figure 2 B.

**Figure 1.**
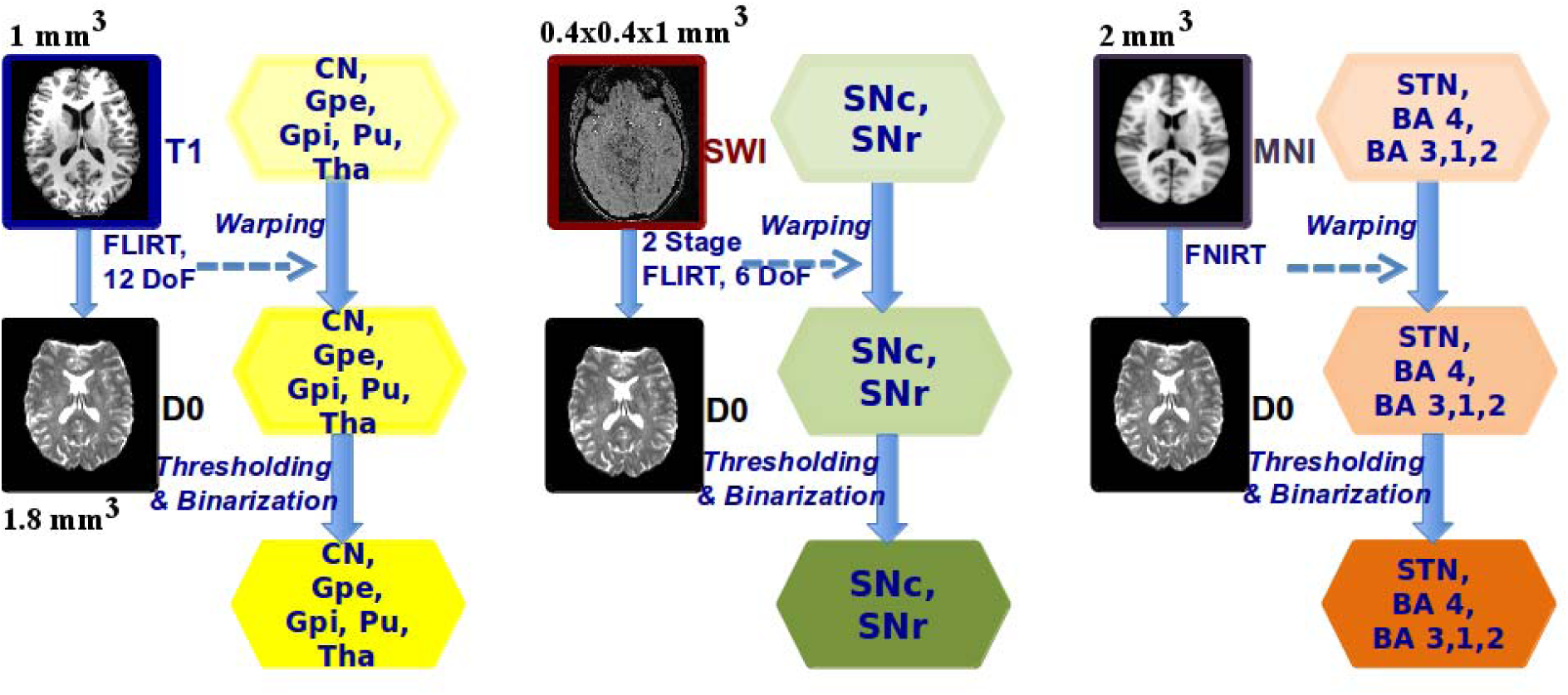
Data processing pipeline steps for manual and atlas based automated segmentation of regions of interest. The modules illustrate alignment of T1-weighted, SWI and MNI atlas images with the non-diffusion-weighted images of the participant. The transformation matrices resulting from each process are used to transform the ROI masks to the diffusion space. The transformation resulting from T1-weighted data alignment was used for CN, GPe, GPi, Pu and Tha (shown in yellow color background), SWI data alignment was used for SNc and SNr (shown in green color background) and MNI atlas data alignment was used for STN, BA4 and BA 3,1,2 (shown in light brown background).

**Figure 2.**
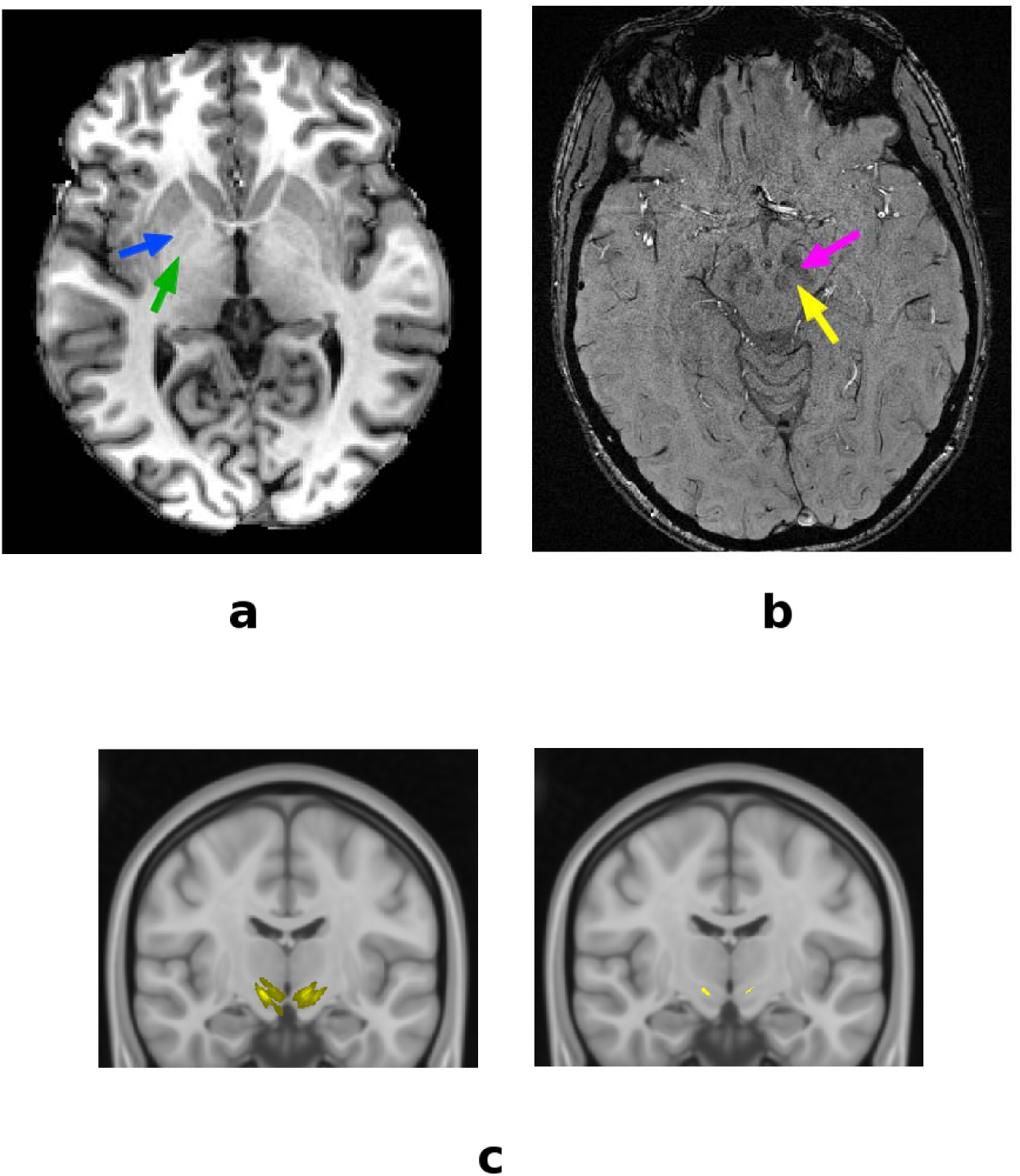
GPe and GPi are segmented based on the T1-weighted images and the separatation between them is shown using blue and green arrows in (a). SNr and SNc are segmented using SWI data as the boundaries of the identifiable area between them is shown in (b) using yellow and pink arrows. The probabilistic STN ROI overlaid on a representative slice from a probabilistic MNI atlas is shown on the left side in (c). The STN mask is thresholded at 30-50% and the thresholded mask is shown on the right side in (c).

In order to identify STN, we referenced a FSL based probabilistic MNI atlas with a resolution of 0.5 ⨯ 0.5 ⨯ 0.5 mm^3^ [Forstmann et al. 2012]. The atlas-based ROI mask was thresholded at 35-50%. This range was chosen for determining the region with the maximum percentage of participant overlap used for creating that atlas [Forstmann et al. 2012]. The STN mask was warped to the MNI atlas with 2 mm^3^ isotropic resolution as shown in Figure 1. Similarly, Broadman Area (BA) 4 (motor cortex) and BA 3,1,2 (somatosensory cortex) were mapped using the probabilistic MNI atlas, with a resolution of 2 ⨯ 2 ⨯ 2 mm^3^. The non-linear transformations between the abovementioned MNI atlases and the non-diffusion-weighted images were calculated using FMRIB’s non-linear registration tool. STN, motor cortex and somatosensory cortex masks were subsequently warped to the diffusion space using these transformations for each participant’s data. Figure 2 C, shows the probabilistic STN ROI overlaid on a representative slice from the probabilistic MNI atlas. The ROI masks of STN and cortex were thresholded with value 0.1 (also see Figure 1).

All binarized ROI masks produced in diffusion space were overlayed on the non-diffusion-weighted image for each participant to perform the last, manual check for sufficient regional coverage. Finally, overlapping mask regions were discarded before ROI masks were used for the probabilistic tractography and the subsequent analysis. Figure 3 A shows an axial view of the subcortical ROI masks for a representative participant, and Figure 3 B and C show coronal and sagittal views of masks for motor cortex (BA 4) and somatosensory cortex (BA 3,1,2), respectively, in addition to the subcortical ROI masks.

**Figure 3.**
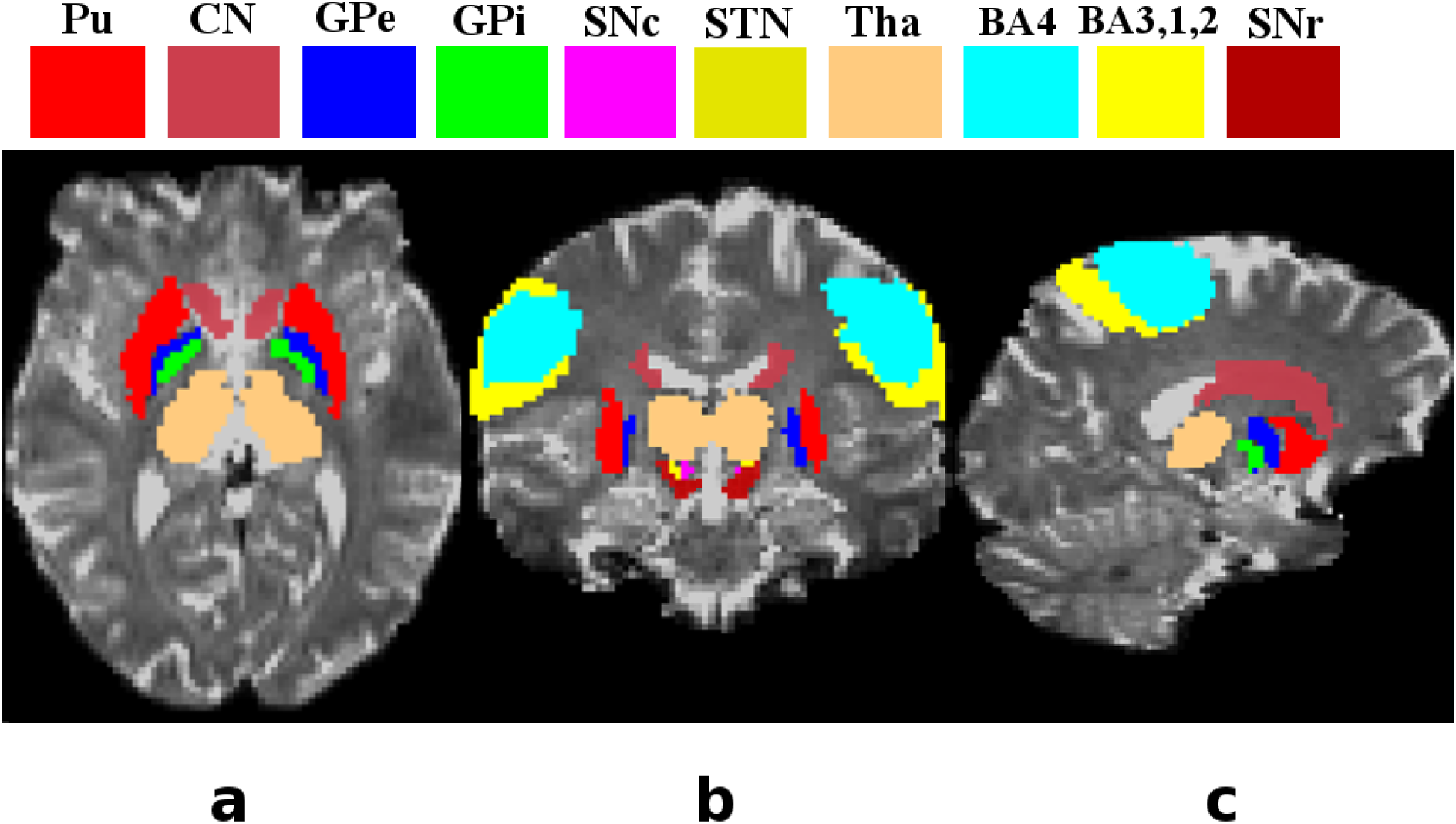
Axial (a), coronal (b) and sagittal (c) views of the subcortical, BA4 and BA 3,1,2 ROIs used in this study. All ROIs are color coded as shown on the top in the figure.

#### Probabilistic tractography

After applying BEDPOSTX, we co-registed the data in diffusion space, structural space and standard space, after which the connectivity distributions of white matter pathways between all subcortical and cortical ROIs were estimated with the probabilistic tractography tool of FDT. The tractography analysis was performed in diffusion space, with 5000 samples and a curvature threshold value of 0.2. The tractography tool was used in ‘classification targets’ mode.

#### Connectivity based seed classification

We performed probabilistic tractography with seed classification in order to gather information about the connectivity patterns between different ROIs within basal ganglia and thalamus; as well as between these regions and motor and sensory cortex. This classification analysis was performed in a voxel-based approach in order to identify the connectivity based topographic subdivisions within each subcortical structure and to quantify the relative frequency of connections between these structures. This approach resulted in structural subdivisions of a specified ROI based on the specific pathways originating from that ROI. Seed classification was performed to identify the connectivity pathways between each seed ROI and the rest of the ROIs that were regarded as targets. This process resulted in a version of a seed ROI mask, called resulting mask, which demonstrated the likelihood of connectivity of each voxel in that ROI with the rest of the subcortical and cortical ROIs. The voxel values of the resulting mask showed the number of tracts originating from the seed ROI and reaching the target ROI successfully. Since the total number of tracts that were generated from each seed voxel was 5000, dividing each voxel value in the resulting mask by 5000 gives the relative frequency of connectivity at that spatial location or voxel.

Subcortical territories were categorized by the voxels of each structure connected to another target structure according to the proportion of connections incepted from the seed region and reaching the target. We quantify the connectivity strength variations of each structure by using the following ranges of the calculated relative frequencies: high range–greater than or equal to 80%, significant range–50-79%, moderate range–10-49% and low range–5-9%. These relative frequency ranges were obtained for each participant and the mean values were calculated across five subjects (i.e. total voxels in the specific range divided by the total number of voxels of the ROI mask). These values were used for quantifying the streamlines connecting each pair of structures of basal ganglia, thalamus and cortical structures. This method did not generate pathways between the subcortical and cortical areas or within the subcortical areas, instead it quantitatively parcellates the brain regions corresponding to the seed ROI masks. Hence, the parcellation of each subcortical area of the basal ganglia and thalamus into distinct subdivisions was performed based on their connectivity profiles with other subcortical and cortical areas.

## Results

### Volumes of regions-of-interest identified in pipeline

In order to ensure the validity of our data processing pipeline, we also estimated the size of all ROIs in the basal ganglia and thalamus for all subjects. Figure 4, shows that our estimated structural volumes were close to those reported in the literature [Anastasi et al., 2006; Harman et al., 1950; Von Bonin et al., 1951; Yelnik, 2002; Chaddock et al., 2010; Ziegler et al., 2013; Menke et al., 2010; Chowdhury et al., 2013; Bagary et al., 2002; Spoletini et al., 2011], with the exception the of STN, which was different from the values documented, for example, in [Forstmann et al., 2012; Massey et al., 2012; Lambert et al., 2012; Nowinski et al., 2005; Colpan et al., 2010]. This may be due to our selective probabilistic thresholding used for STN localization, which is different from previous methods [Lambert et al., 2012], where STN was localized as one large region containing iron. The STN volume estimated by us (64.1 mm^3^ in the case of whole brain coverage) is closer to the STN volume, which is typically used for the deep brain stimulation.

**Figure 4.**
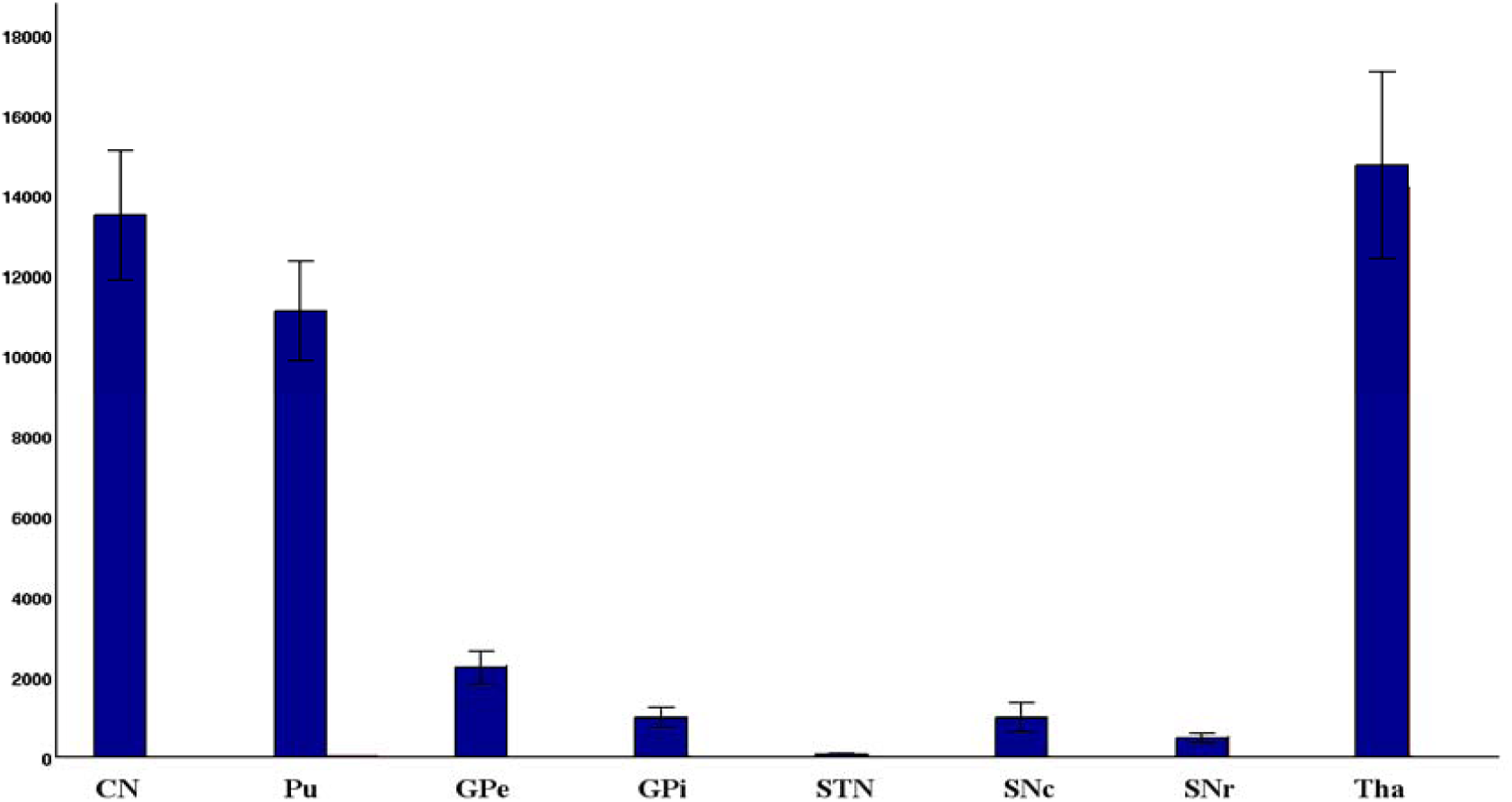
Volumes of the subcortical regions used in this study are shown in this figure. The volume of each region is averaged across five participants as illustrated in figure.

### Probabilistic tractography in pipeline

Figures 5, 6 and 7, show for each subcortical structure the mean percentages for high, significant and moderate -strength connections, respectively (except in the case of the STN region where the low strength range is also used). Corresponding connectivity patterns for one representative participant are shown in Supplemental Figures 1-8. We next describe the obtained connectivity patterns for each structure.

**Figure 5.**
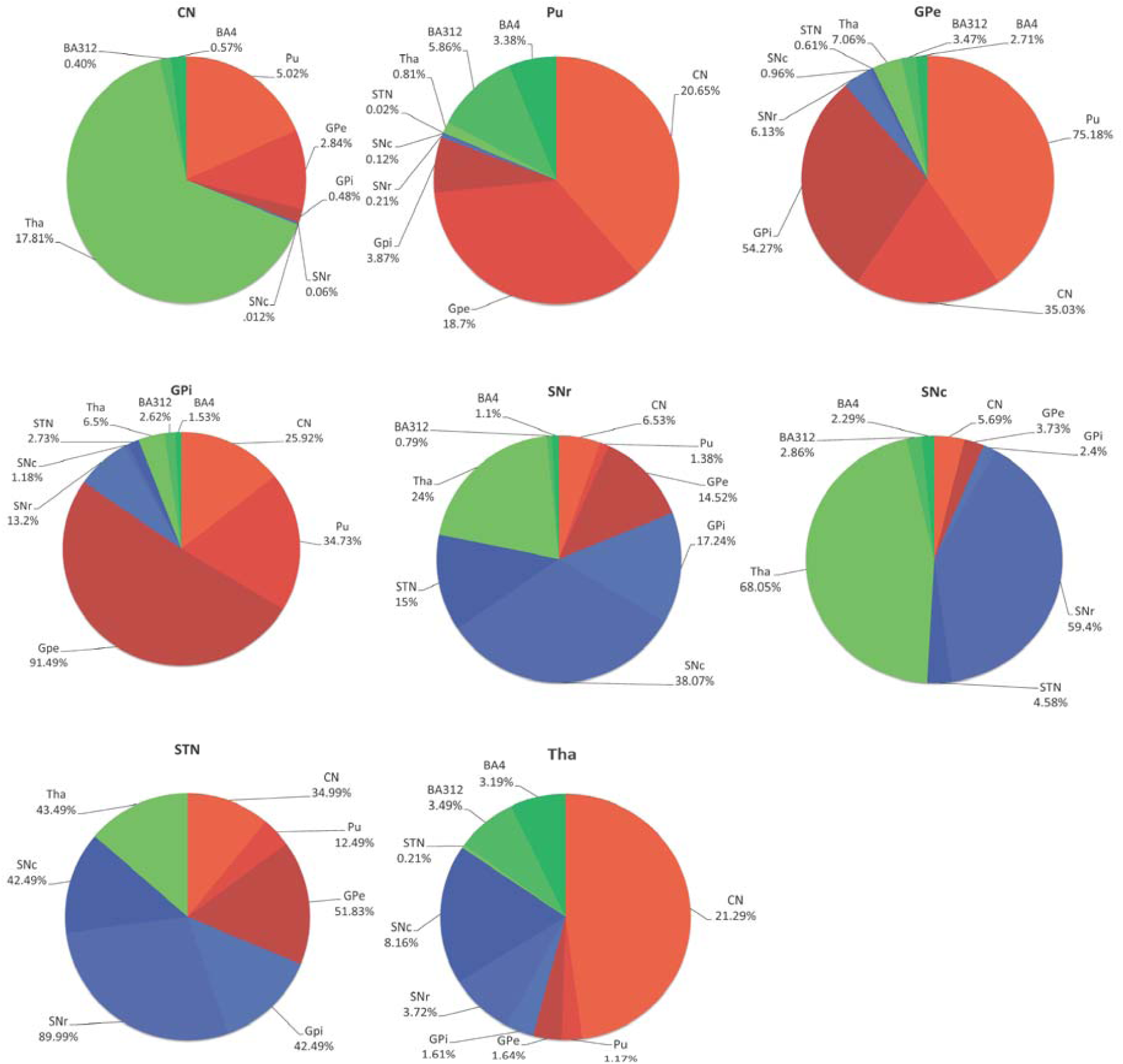
“High range” connectivity patterns. Percentages of the connections that were catagorized in the ‘high range’ (as described in the methods section) between source and target subcortical regions are calculated with respect to the total connections originating from source subcortical region. CN Caudate Nucleus, Pu Putamen, GPe Globus Pallidus External, GPi Globus Pallidus Internal, SNr Substantia nigra pars reticulata, SNc Substantia nigra pars compacta, SN Subthalamic nucleus, Tha Thalamus

**Figure 6.**
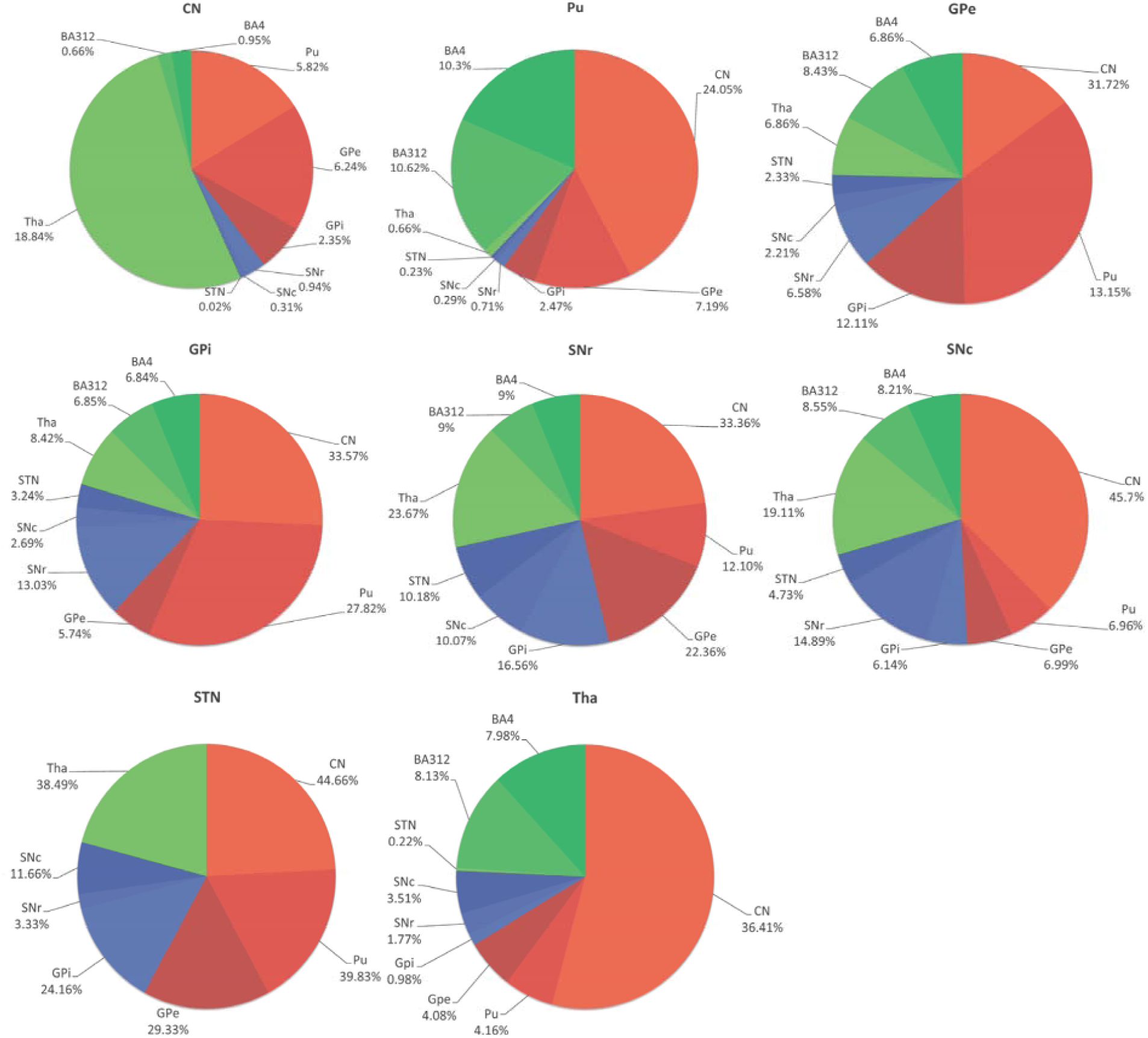
“Significant range” connectivity patterns. Percentages of the connections that were catagorized in the ‘sigificant range’ (as described in the methods section) between source and target subcortical regions are calculated with respect to the total connections originating from source subcortical region. CN Caudate Nucleus, Pu Putamen, GPe Globus Pallidus External, GPi Globus Pallidus Internal, SNr Substantia nigra pars reticulata, SNc Substantia nigra pars compacta, SN Subthalamic nucleus, Tha Thalamus

**Figure 7.**
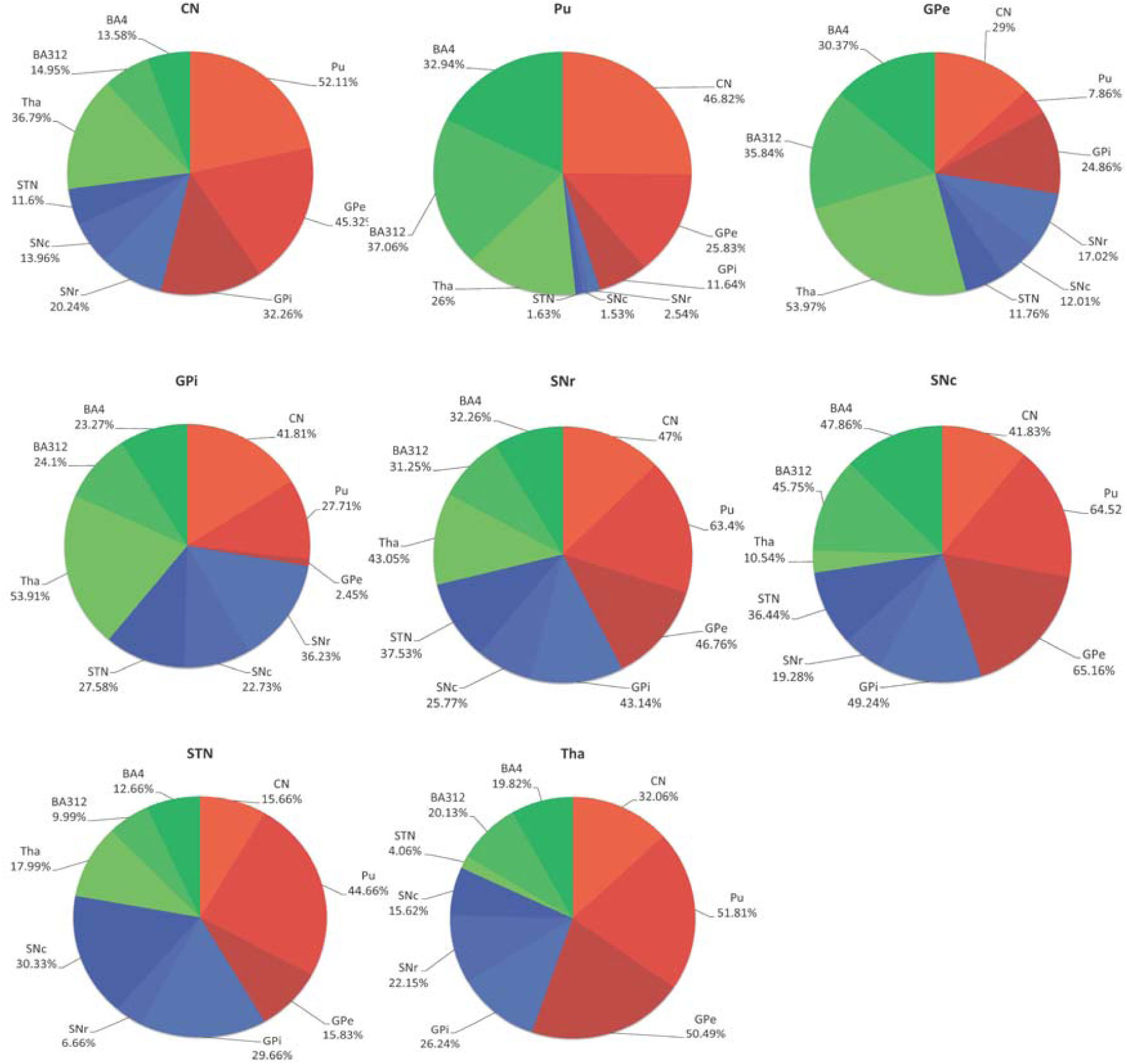
“Moderate range” connectivity patterns. Percentages of the connections that were catagorized in the ‘moderate range’ (as described in the methods section) between source and target subcortical regions are calculated with respect to the total connections originating from source subcortical region. CN Caudate Nucleus, Pu Putamen, GPe Globus Pallidus External, GPi Globus Pallidus Internal, SNr Substantia nigra pars reticulata, SNc Substantia nigra pars compacta, SN Subthalamic nucleus, Tha Thalamus

#### Caudate Nucleus (CN)

CN was strongly connected to Pu, GPe and Tha. Some connections between CN and GPe had a higher range of relative frequency. Connections between CN and GPi were in the significant relative frequency range. CN was connected to SNc and STN through a moderate range of relative frequency of pathways, whereas it was connected to SNr through significant range of relative frequency of projections. The detailed explanation and illustration of CN connections are given in Supplemental Figure 1.

#### Putamen (Pu)

Pu had strong connections with CN, GPe and GPi. Moderate relative frequency connections were found originating from CN and terminating in SNr, SNc (only a few) and Tha. Supplemental Figure 2 shows the connectivity pathways of Pu and its corresponding relative frequency.

#### Globus pallidus external (GPe) and internal (GPi)

GPe was strongly connected to Pu and GPi, anterior and medial portion of GPe contained high relative frequency connections to CN, SNr and SNc. Although very few voxels of rostral and ventro-caudal GPe region were found that had high relative frequency of connectivity with STN. Dorsal and lateral GPe was connected with Tha in a similar fashion. Supplemental Figure 3 illustrates the projections originating from GPe.

GPi had substantial high relative frequency connections with GPe, CN, Pu and SNr. Rostral GPi contained very few strongly connected voxels with STN, whereas dorsal and lateral portions of GPi had very few significant relative frequency projections with Tha. Only a few voxels of GPi were strongly connected with SNc. A detailed explanation and illustrations of GPi connectivity with other ROIs are given in Supplemental Figure 4.

#### Substantia nigra pars compacta (SNc) and pars reticulata (SNr)

Due to close proximity, SNr and SNc were strongly connected to each other. A few voxels of lateral and medial portions of ventral SNc comprised projections to CN and Pu had high and significant relative frequencies, repectively.

Ventral SNc and ventral-ipsilateral SNc were connected to GPe through moderate relative frequency connections and to GPi through significant relative frequency connections, respectively. Medial SNc had strong connections to STN but it was comprised of very few voxels, whereas the connections from the posteriolateral SNc to Tha were in the low range.

Lateral ventral SNr was strongly connected to CN, GPe and GPi. It also had connections with Pu in the significant relative frequency range. Anterior medial SNr was primarily connected to CN. Medial SNr was highly connected to STN and Tha, whereas posterior SNr was also strongly connected to Tha. The detailed explanation and illustrations of SNc and SNr projections are given in Supplemental Figures 5 and 6, respectively.

#### Subthalamic nucleus (STN)

Since deep STN region is targeted, the size of the seed ROI is very small. Subsequently, the low range connections were found that originated from STN and terminated in the rest of the subcortical areas, as illustrated and described in Supplemental Figure 7.

#### Thalamus (Tha)

Tha comprised strong connections with CN and STN. It contained projections to Pu that had significant and moderate connectivity relative frequencies. Tha was connected with GPe, GPi, SNr and SNc through pathways of high and significant range connectivity, as demonstrated and explained in Supplemental Figure 8.

## Discussion

We used a multiband diffusion imaging protocol and processed the acquired data with a pipeline designed for performing probabilistic tractography in order to obtain a detailed reconstruction of the white matter tracts that comprise the basal ganglia and thalamus. Such connectivity maps could play an increasing role in retrieving image-based prognostic information from data of patients with basal ganglia pathology, or with regard to movement disorder treatment evaluation. Our data processing pipeline can be easily set up in a clinical setting, with an MRI field strength of 3T. We next discuss our results and provide, where possible, a comparison with the previous literature on basal ganglia connectivity.

### Caudate Nucleus (CN)

CN is considered as the signal receiving part of the basal ganglia. As Figure 5 and Supplementary Figure 1 illustrate, we found strong connections that emerge from ventral and dorsal CN to Pu. We also located specific origins of the projections from CN to both GPe and GPi and vice versa. We found CN to be strongly connected to the dorsomedial GPe [Lenglet et al., 2012] and ventromedial GPi, whereas dorsolateral portions of CN were well connected to both SNr and SNc (Lynd-Balta and Haber, 1994; Lynd-Balta and Haber, 1994, also see Supplemental Figure 1). The medioventral CN was connected to the deep STN [Smith and Parent, 1986; Parent and Smith, 1987], whereas the the dorsolateral and rostroventral CN had strong projections to Tha. There were also corticostriatal projections from BA 4 and BA 3,1,2 to the dorsoposterior sections of CN [Draganski et al., 2008; Flaherty and Graybiel 1991; Flaherty and Graybiel 1993]. Our findings are well in agreement with previous work on CN connectivity [e.g. Parent and Hazrati, 1995; Alexander and Delong, 1985; Gerfen et al., 1996; Wilson, 1998; Smith et al. 1998].

### Putamen (Pu)

Pu receives input through the excitatory glutamatergic projections from the cerebral cortex and thalamus, modulatory dopaminergic inputs from the SNc and ventral tegmental area (VTA), and serotonergic inputs from the dorsal raphe nucleus (DR). Pu also connects directly, and indirectly via GPe and STN, to the output nuclei of the basal ganglia i.e. GPi and SNr. We found strong intrastriatal connections that arise from Pu to CN as can be seen in Figure 5 and Supplementary Figure 2. The straitopallidal connectivity described in [Draganski et al., 2008] implies that the Pu connections occupied mainly the ventrolateral pallidum and the dorsolateral Pu is connected with both pallidum [Francois et al., 1994; Parent et al., 2000; Saleem et al., 2002] and premotor cortical area. However, we found that efferents from Pu to GPe [Lenglet et al., 2012] were primarily located in medial Pu, whereas a few efferents were found from medial, ventral and dorsolateral Pu to GPi. A medio-laterally decreasing gradient of projections from Pu to SN were identified by Lenglet et al. [2012]. We could not find significant connections between Pu and SN as the relative frequencies of connections was found to be very low for this pathway. STN neurons that excite Pu are located in the sensorimotor dorsolateral two-thirds of STN [Smith and Parent, 1986; Parent and Smith, 1987]. Our results are based on the deeply localized STN; therefore, we could not find significant projections between Pu and STN. Most ventral parts of Pu are connected with both Tha and prefrontal and orbitofrontal cortex, [Draganski et al., 2008]. It was shown by Draganski et al. [2008] that the central Pu region connects with ventral and dorsal premotor cortical areas and overlaps more ventrally with dorsolateral prefrontal cortex and caudally with motor area M1-associated connections. There are convergent projections from frontal cortical motor areas and some interconnected ventral thalamic motor relay nuclei to large territories of the postcommissural Pu [McFarland and Haber, 2000]. Our results indicate that Pu contains high, significant and moderate connectivity pathways to BA 4 and BA 3,1,2. We observed that the ventral region (both anterior and posterior) of Pu contains projections to Tha and this region overlaps more ventrally with the region that is primarily connected to cortical areas BA 4 and BA 3,1,2. (Supplemental Figure 2).

### Globus pallidus external (GPe) and internal (GPi)

GP internal (GPi) and GP external (GPe) are separated by the lamina pallidi medialis. CN and Pu transmit signals to GPi via indirect and direct pathways. The indirect pathway is composed of the CN and Pu neurons that project to GPe in the first phase. Thereafter, GPe sends signals to STN, which carries them to GPi and SNr. Apart from that, GPe initiates GABAergic projections to GPi, SNr and the reticular thalamic nucleus [Parent and Hazrati, 1995; Smith et al., 1998], and projection from the GPe to the CN and Pu regions also exists [Bevan et al., 1998]. We found that the medial GPe is strongly connected to GPi, whereas a portion of the ventral GPe is connected to SNc, as shown in Supplemental Figure 3.

CN is strongly connected to the dorso-medial third of GPe, and Pu mainly projects to the ventrolateral two-thirds of GPe [Lenglet et al. 2012]. Our results are in agreement with these previous findings i.e., the lateral GPe is primarily connected to the medial portion of Pu and its rostrodorsal portion is connected to CN. The most rostroventral portion of GPe was found to be connected to Tha and STN [Lenglet et al. 2012]. The interconnected neurons of GPe and STN innervate, via axon collaterals, functionally related neurons in GPi [Shink et al. 1996] and therefore, populations of neurons within the sensorimotor, cognitive, and limbic territories in GPe are reciprocally connected with populations of neurons in the same functional territories of STN. In turn, neurons in each of these regions innervate the same functional region of GPi. We found some projections originating from the rostroventral and ventromedial GPe to Tha and STN, which are illustrated in Supplemental Figure 3. The strong connectivity between inferior-posterior GPe and BA 4 and BA 3,1,2 is also verified by our results.

Our results further show that the latero-caudal GPi is connected to Pu, the rostral GPi is mainly connected to CN and the central GPi is strongly connected to GPe. The mid-portion of GPi is connected to SN, whereas its rostral portion is divided into a dorsal part connected to CN, and a ventral part connected to STN and Tha [Lenglet et al. 2012]. The pallidothalamic projection stems from the ventral anterior/ventral lateral (VA/VL) thalamic nuclei [Parent and Hazrati, 1995; Sidibe et al., 1997]. However, we found that ventral and medial GPi is mainly connected to SNc, whereas a minor portion of the inferior-posterior GPi connected to SNr, STN and Tha also exists. These connectivities are illustrated in Supplemental Figure 4. [Draganski et al. 2008] found projections from pallidum to sensorimotor, prefrontal, orbitofrontal and premotor cortical areas. In comparison to those results, our study is more specific as connectivity analysis is done separately for GPe and GPi instead of single pallidum region (combined GPe and GPi). We found strong central GPi connections with BA 4 and BA 3,1,2 cortical regions that can be seen in Supplemental Figure 4. Several findings regarding connectivity of GPi and other subcortical nuclei exist in literature [Simth et al. 1998; Baron et al. 2001; Parent and DeBellefeuille 1982] which correlate with our results. However, they employed histological and staining methods which are different in terms of specificity from our computer based method.

### Substantia nigra pars compacta (SNc) and pars reticulata (SNr)

. The dendrites of individual SNr neurons largely conform to the geometry of striatonigral projections, and the substantial primary axons of SNc project to the ipsilateral GP [Mettler 1970]. These connectivity pathways of SNc and SNr are illustrated in Supplemental Figure 5 and 6, respectively. It can be seen in our results that the lateral and anterior SNr project to CN and Pu. It was found by Lenglet et al. [2012] that projections from SN to GPi are mainly ventral, whereas nigrostriatal fibers project primarily to Pu medially, dorsally and ventrally. We found that the projections from SNr to GPe and GPi are primarily ventral and lateral, whereas the projections from SNc to GPe and GPi are ventral and ipsilateral. It was also demonstrated by Lenglet et al. [2012] that the most anterolateral portion of SN was strongly connected to GPi, GPe and striatum while the posterolateral portion was strongly connected to Tha. It was hypothesized in [Lenglet et al., 2012] that the medial SN, -well connected to GP but not to the striatum - comprises part of the SNc. Our results are consistent with this hypothesis since we find the medial SNc region to be strongly connected to GPe and GPi, but not well connected to Pu. SNr is one of the main projection sites of the STN [Parent and Smith, 1987] and the STN projections to SNc are additional indirect pathways through which signal from cortical regions reaches basal ganglia output structures. SNc-STN and SNr-STN pathways depicted in Figure 5 and 6, respectively, show that medial regions of SNc and SNr are strongly connected with STN. As the resolution of STN region used in our work is low, it was not possible to parcellate the pathways accroding to connections with STN sub-territories. Thalamic connections originating from the medial part of SNr terminate mostly in the medial magnocellular division of the VA (VAmc) and the mediodorsal nucleus (MDmc) [Ilinsky et al., 1985]. The lateral part of the SNr contains projections to the lateral posterior region of the VAmc and to different parts of MD. Connections between SNr and rostral and caudal intralaminar thalamic nuclei [Ilinsky et al., 1985; Mailly et al., 2001], and the nigro-intralaminar thalamic projection ending in the PF area [Smith et al., 2000] also exist. Connections from SN to thalamic nuclei, such as the medio-dorsal nucleus (MD), ventrocaudal nucleus (Vc), latero-polar nucleus (Lpo) and ventro-oral anterior nucleus (Voa) were demonstrated in [Lenglet et al., 2012]. Our results are in agreement with such previous findings as strong connectivity strength of SNc-Tha pathways is evident in Supplemental Figure 5 and the similar pattern for SNr-Tha connections can be seen in Supplemetal Figure 6. Our results demonstrate that central SNr is well connected to the BA 4 and BA 3,1,2 areas; and the inferior and posterior SNc are well-connected to these cortical regions.

### Subthalamic nucleus (STN)

The STN is considered as the minor receptor of the basal ganglia because STN receives excitatory glutamatergic inputs from the cerebral cortex and thalamus, modulatory dopaminergic inputs from the SNc and ventral tegmental area (VTA), and serotonergic inputs from the dorsal raphe nucleus (DR). The main projection sites of STN are GPe, GPi, and SNr. The STN projections to SNc, CN and Pu are additional indirect pathways through which the information is transferred to the basal ganglia. Particularly, the STN is comprised of segregated sensorimotor, associative, and limbic territories. In our work, due to very small size of localized STN, it was not possible to comment on the connectivity properties of the sub-territories of STN as specified in [Lambert et al. 2012]. Supplemental Figure 7 illustrates our findings related to STN connectivitiy.

### Thalamus (Tha)

One of the main sources of glutamatergic inputs to striatum is the thalamo-CN projection, which originates mainly from the centromedian (CM) and parafasicular (PF) intralaminar nuclei of Tha. The CM projects to the sensorimotor territory of CN and Pu, whereas the PF projects mainly to the associative territory and slightly to the limbic territory of CN and Pu [Sadikot et al., 1992; Sidibe and Smith, 1996]. Our results give evidence of existence of these thalamic projections as we notice that the lateral Tha projects mainly to the dorsal CN area in Supplemental Figure 8. The input to the limbic territory of CN and Pu emerges primarily from midline and rostral intralaminar nuclei of Tha, which can be observed in Supplemental Figure 8. Although associative region of Tha also projects to CN and Pu, but less than intralaminar nuclei [Groenewegen and Berendse, 1994; Giménez-Amaya et al., 1995]. Connections from the medial thalamic nuclei to ventral striatal areas and the lateral thalamic nuclei to dorsal striatal areas were found in [Draganski et al., 2008]. Projections from thalamic CM/PF, medio-dorsal nucleus (MD), latero-polar nucleus (Lpo), ventro-oral anterior nucleus (Voa) and the dorso-intermediate nucleus (Dim) to CN were demonstrated in [Lengelt et al., 2012]. These findings are in agreement with our thalamo-CN connectivity results shown in Supplemental Figure 8. In case of thalamo-Pu projections, the precommissural Pu receives inputs from the dorsolateral PF area [Sidibe et al., 2002]. Our results illustrated in Supplement Figure 8 agree with these findings. Most of the thalamo-Pu connections found in our work exist in the moderate relative frequency range, especially in the CM and PF regions. Contrary to that, the thalamic region having strong connections comprises pulvinar, ventral lateral and ventral posterior nuclei. Thalamic connections with ventromedial pallidum were shown in [Draganski et al., 2008]. The central GPi’s pallidal neurons that project to thalamic motor nuclei send axon collaterals to the caudal intralaminar nuclei [Parent et al. 1999]. Major associative inputs from the GPi end in the dorsolateral extension of PF (PFdl) thalamic region, which does not project back to CN but rather connects with the precommissural putamen. The limbic GPi is connected with the rostrodorsal part of PF, which projects back to the nucleus accumbens [Giménez-Amaya et al., 1995]. Ventral MD of Tha is connected to ventral GP and vice versa [Lengelt et al., 2012]. In particular, connectivity between caudal to rostral Tha and GPi and medial GPe was demonstrated in [Lengelt et al., 2012]. Moreover, it was shown that CM and Lpo are connected with GPi [Lengelt et al., 2012]. Our results are in compliance with these previous findings. The connectivity pathways from rostral ventral anterior and MD nuclei to GPe and GPi are verified by our results in Supplemental Figure 8. The nigrothalamic connections from SN to different Tha nuclei, such as ventrocaudal nucleus (Vc), MD, Lpo and Voa have been identified [Lengelt et al., 2012]. Comparatively, our results are more specific as demonstrated by the strong connectivity of SNr with Vc, MD, Voa; and SNc with Vc and Voa. It was shown previously that GPi and SNr are connected to ventro-intermediate (Vim), Voa, Lpo; additionally vim projects to STN [Lengelt et al., 2012]. We found similar projections, although the range of relative frequencies of connectivities is between significant and high. Rostral ventral anterior and MD nuclei project to both pallidum and dorsolateral prefrontal cortex [Draganski et al., 2008]. Ventral lateral Tha projects to pallidum; Pu and motor area; and premotor area [Draganski et al., 2008]. We found that the similar region of Tha has connections with Pu and BA 4 in the high relative frequency range and with GPe in the significant relative frequency range. Projections from the caudal dorsal Tha to somatosensory cortex are also demonstrated in Supplemental Figure 8.

Taken together, these results show overall good agreement with the previous literature, thus, our pipeline proves to be suitable for performing detailed topographic analysis of subcortical structural connectivity patterns found in the human brain. This pipeline is ideal for processing imaging data acquired during experiments specifically designed for enhanced basic research, moreover it also has great potential for application in disease identification or treatment evaluation for a range of movement, neuropsychiatric, or neuropsychological disorders.

### Limitations

As with any imaging technique, various sources of image distortion could hamper the quantification of probabilistic connectivity patterns using diffusion MRI. More specifically, motion, eddy-current, susceptibility, main magnetic field inhomogeneity, concomitant fields and any other types of distortions must be precisely corrected in the data processing pipeline in order to ensure improved quantitative processing of the multi-contrast MRI data. Due to the imperfect distortion correction, distant proximity of certain structures and dense crossing fibers, the relative frequencies of connectivity of some structures were not in the “high range” in our work, for example between STN and cortical areas, CN and STN, Pu and STN, SN and cortical areas, SN and Pu, etc. However, it is important to note that this pipeline used data acquired at 3T, where visibility of STN using routine clinical MRI is limited. Therefore, specifically optimized T_2_* sequences can be employed in the protocol for improved visualization of STN to make accurate identification and delineation of the nucleus and target the motor portion of STN. As connections revealed through diffusion MRI represent the sum total of both afferents and efferents, the inability of determining the polarity of connections may be another limiting aspect of this study.

## Conclusions

We demonstrated the working principles of a data processing pipeline specifically designed for the structural multiband MRI data. This data processing pipeline produced results which show good agreement with previous research on basal ganglia connectivity. The designed pipeline confirms that the high resolution, combined with high SNR per unit time of acquisition, afforded by the multiband diffusion MRI sequence can be beneficial for the identification of subcortical connectivity patterns. The taxonomy of connectivity maps resulted by the proposed pipeline, when generated using large patient cohort, might have a potential role in retrieving image-based prognostic or therapeutic information of patients with basal ganglia pathology. Our strategy, which is based at 3T field strength, can thus be adapted to any relevant clinical environment as well as by the broader brain research community.

## Supporting information

Supplement 1 (CN Connectivity Map)

Supplement 2 (Pu Connectivity Map)

Supplement 3 (GPe Connectivity Map)

Supplement 4 (GPi Connectivity Map)

Supplement 5 (SNc Connectivity Map)

Supplement 6 (SNr Connectivity Map)

Supplement 7 (SNc Connectivity Map)

Supplement 8 (Tha Connectivity Map)

