## Supplement Text for "Data Processing Pipeline for Multiband Diffusion-, T1- and Susceptibility Weighted MRI to Establish Structural Connectivity of the Human Basal Ganglia and Thalamus"

**Caudate Nucleus (CN)**

In the context of connectivity between CN and Pu, the anterior regions of CN had highest relative frequencies (greater than 80%) of connectivity in the inferior slices and the posterior regions had shown highest relative frequencies (greater than 80%) in the superior slices, as shown in the axial section of Supplement 1 A. In the coronal section of CN shown in Supplement 1 A, the superior regions had shown highest relative frequencies of connectivity with Pu as the slices are traversed from anterior to posterior. In the saggital view of Supplement 1 A, the highest relative frequencies were found in the head, corpus and tail of CN. In case of connectivity between CN and GPe, lateral parts of CN had shown higher relative frequencies of connectivity (greater than 80%), as shown in the axial view in Supplement 1 B, while medial parts showed significant connectivities (50-79%) as slices are traversed from lowest to highest position. The main connections from CN to GPi originated from the region having significant relative frequencies (70-80%) within the CN’s head and they were localized more centrally as shown in Supplement 1 C. Overall, moderate relative frequencies of connectivity (10-49%) were observed between CN and SNc, where most of the moderate relative frequency connections (49%) were found in the head and medial region of CN as illustrated in Supplement 1 D. In case of connectivity between CN and SNr, significant relative frequencies (50-70%) were widespread as shown in Supplement 1 E. It is evident in Supplement 1 F that ventral part of CN’s head contained connections to STN having moderate relative frequency (25-30%), whereas the tail of CN showed low relative frequencies (5-15%) of connectivity. It can be observed in Supplement 1 G that highest relative frequency voxels of CN connected with Tha (grater than 80%) were found along the posterior and lateral portions of CN. Highest number of voxels of CN, which had highest connectivity relative frequencies (greater than 80%), were connected with Tha. The projections emerging from CN to cortical areas BA 4 and BA 3,1,2, with highest relative frequency of connectivity (greater than 90%), were localized in more dorsoposterior region of CN, especially in tail. Nevertheless, these connections were less in number as shown in Supplement 1 H and I.

**Putamen (Pu)**

Tractography resulted in very strong projections from Pu to CN with highest relative frequencies of connections (greater than 80%), as it can be seen in Supplement 2 A. High relative frequencies (greater than 80%) were found in the case of connections between Pu and GPe, as shown in Supplement 2 B; but less number of connected voxels were resulted compared with the connected voxels between Pu and CN. In case of connectivity between Pu and GPi, very few high relative frequency connections (greater than 80%) were observed near the midline region as shown in axial view of Supplement 2 C. A few connections originating from Pu to SNc and SNr were found as shown in Supplement 2 D and E, respectively. The relative frequencies of connections with SNr (50%) were slightly higher than the relative frequencies of connections with SNc (30%). The anterior and posterior regions of ventral Pu showed moderate relative frequencies of connectivity (around 30%) with Tha as illustrated in Supplement 2 F. In the context of connections of Pu with BA 4 and BA 3,1,2, posterior regions of Pu showed high relative frequencies of connections with both cortical areas (90%) as shown in Supplement 2 G and H.

**Globus pallidus external (GPe) and internal (GPi)**

It is illustrated in Supplement 3 A that most of the anterior and medial portion of GPe was strongly connected (relative frequency range 80-90%) with CN as slices are traversed from ventral to dorsal. Similarly, GPe had highest relative frequencies of connectivity (greater than 90%) with both Pu and GPi as shown in Supplement 3 B and C, respectively. Anterior and medial parts of GPe are primarily connected with SNr (greater than 90%), whereas only a few voxels of GPe appeared to be connected with SNc having highest relative frequencies (greater than 90%) that are shown in Supplement 3 E and D, respectively. Supplement 3 F and G demonstrates that very less high relative frequency (greater than 80%) voxels of GPe were connected to STN and Tha. Rostral and ventro-caudal portion of GPe was found to connect with STN. Dorsal and lateral portions of GPe were connected with Tha. In case of projections originating from GPe to BA 4 and BA 3,1,2, posterior region of GPe had shown highest relative frequencies (greater than 90%), which can be seen in Supplement 3 H and I.

GPi demonstrated strongest connections (greater than 90%) with CN and Pu, as shown in Supplement 4 A and B. GPi and GPe were strongly connected that is evident in Supplement 4 C. Significant region of GPi was connected to SNr with highest relative frequencies (90%) as illustrated in Supplement 4 E. Rostral GPi showed only a few voxels that were connected with STN. Dorsal and lateral portions of GPi were connected with Tha. The GPi’s regions connected to SNc, STN and Tha were composed of very few voxels having relative frequencies of connectivity in the ranges of 90%, 90% and 70%, respectively. Strong connectivity of GPi with BA 4 and BA 3,1,2 was observed and most of the central portion of GPi contained voxels with high relative frequencies of connectivity (greater than 80%), as shown in Supplement 4 H and I.

**Substantia nigra pars compacta (SNc) and pars reticulata (SNr)**

Lateral and medial portions of ventral SNc were well-connected to CN, but only a few voxels of SNc having high relative frequencies were connected to both CN (80-90%) and Pu (70%), as shown in Supplement 5 A and B, respectively. Ventral SNc was connected to GPe with moderate (10-49%) relative frequencies and ventral and ipsilateral SNc projected to GPi with significant (60-70%) relative frequencies as illustrated in Supplement 5 C and D. The voxels of SNc in the proximity of SNr were strongly connected (90%) with SNr that can be seen in Supplement 5 E. Medial SNc was connected to STN where a few voxels were found that had high connectivity relative frequency range (greater than 80%). Posteriolateral SNc region was primarily connected to Tha and it can be observed in Supplement 5 G that very few voxels of SNc were connected to Tha with low relative frequency of connectivity. Inferior part of SNc contained projections to BA 4 and BA 3,1,2 having high range of relative frequency of connectivity (greater than 80%).

Strong connections (relative frequency range 80-90%) between SNr and CN were found in the lateral portion of ventral SNr and also in the anterior medial portion of SNr as shown in Supplement 6 A. Lateral portion of ventral SNr also contained significant connections (70%) with Pu and strong connections (90%) with GPe and GPi, as illustrated in Supplement 6 B and C. Medial portion of SNr was strongly connected (90%) to SNc as shown in Supplement 6 E. Strong projections (90%) from medial SNr to STN were found as illustrated in Supplement 6 F. Posterior and medial portion of SNr was primarily connected (90%) with Tha shown in Supplement 6 G. Significant number of voxels in the medial SNr, shown in Supplement 6 H and I, were found connected with BA 4 and BA 3,1,2 with high relative frequency of connectivity (80%).

**Subthalamic nucleus (STN)**

The STN region localized in this work was very small in size, therefore it was not possible to segregate sub-regions of STN that were connected with other areas. Overall, the relative frequencies of connectivity of projections originating from the STN localized in this work to different regions are shown in Supplement 7. It is important to mention here that some connectivity relative frequencies begin from the initial low range of 5% in the case of STN.

**Thalamus (Tha)**

The representative topological subdivisions of Tha based on the connectivities to other subcortical and cortical regions are shown in Supplement 8. The cytoarchitectonic architecture of Tha is very complex, which is exemplified by different efferents and afferents pertaining to many segregated regions. It can be seen in Supplement 8 A that anterior dorsal, lateral dorsal, medial dorsal, ventral anterior, lateral posterior and pulvinar nuclei of Tha were strongly connected (80%) to CN. Ventral posterior and lateral posterior nuclei were connected to Pu with significant relative frequencies (70%). Medial dorsal and pulvinar were connected to Pu with moderate relative frequencies (10-49%). Anterior dorsal was found to connect to Pu with significant relative frequencies (70%) though a few voxels were found. Medial Tha was connected with ventral CN and Pu regions, whereas lateral Tha was connected with dorsal CN and Pu. Projections from ventral posterior, ventral lateral, ventral anterior, lateral posterior and pulvinar were found to connect to GPe with significant connectivity relative frequencies (50-79%), whereas medial dorsal, anterial dorsal and lateral dorsal were connected to Pu with high relative frequencies (greater than 80%). Large portion of dorsal Tha was connected to GPi with significant relative frequencies (10-49%) and minor portion of ventral anterior Tha was connected with GPi with high relative frequencies (greater than 80%). Ventral anterior, ventral lateral, ventral posterior nuclei were connected to SNc and SNr with high relative frequencies (80-90%) and few voxels in the medial Tha portion were connected to SNc and SNr with significant relative frequencies (10-49%). Lateral posterior and pulvinar were also connected to SNr with significant relative frequencies (10-49%). Ventro-intermediate nucleus (Vim), the main target in Tha for deep brain stimulation, was strongly connected (90%) with STN. These observations manifest high degree of connectivity of Vim with the very deep nuclei i.e., SNc, SNr and STN. Central portion of Tha contained a wide range of voxels having high relative frequency of connectivity (90%) to BA 4 and BA 3,1,2. It is important to mention that low degree of interhemispheric differences were observed in this case. Ventral posterior and ventral lateral portion of Tha was well-connected to the BA 3,1,2 i.e. the somatosensory cortex area.
