## Supplementary figures and images for "Data Processing Pipeline for Multiband Diffusion-, T1- and Susceptibility Weighted MRI to Establish Structural Connectivity of the Human Basal Ganglia and Thalamus"

### Supplement 1 (CN Connectivity Map)

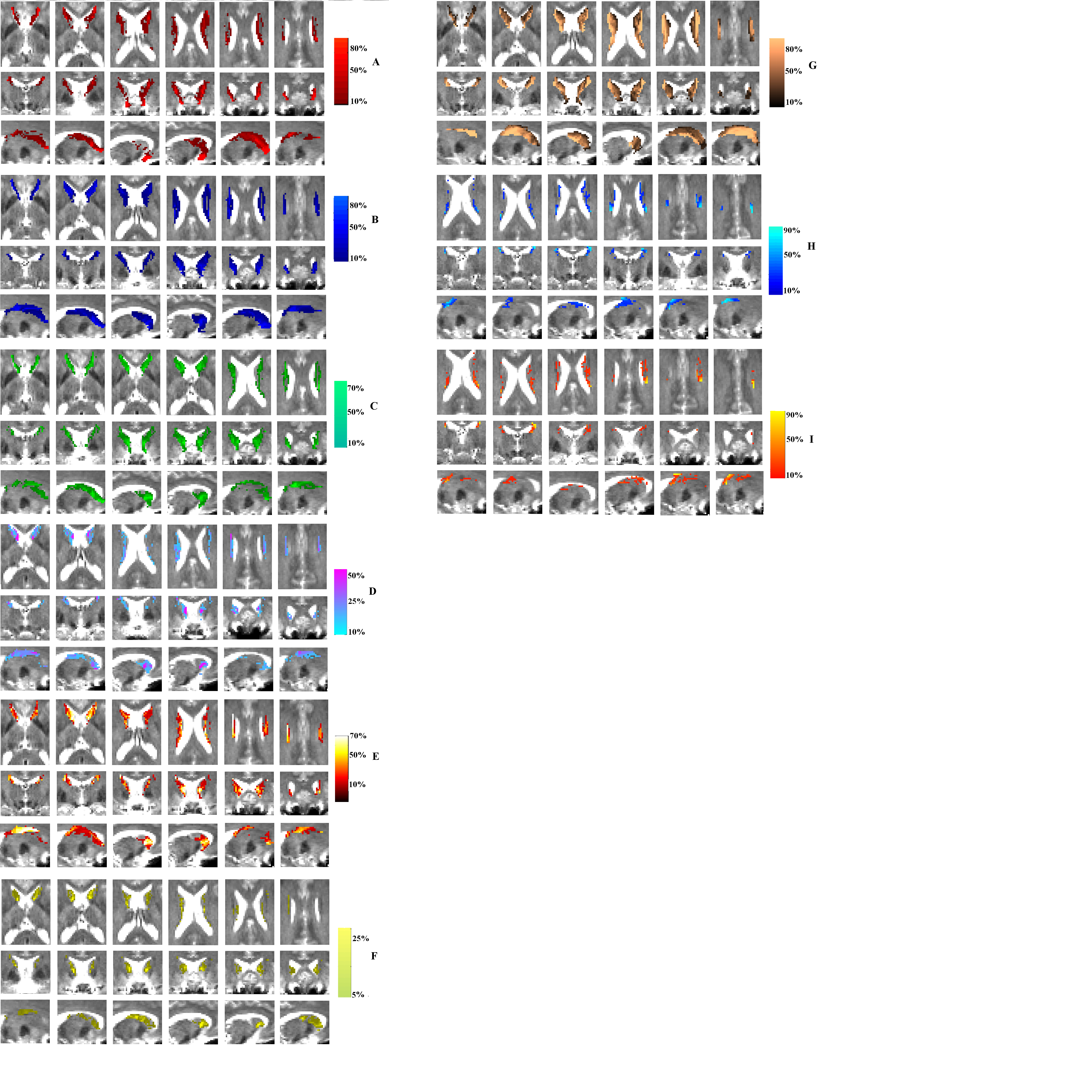

### Supplement 2 (Pu Connectivity Map)

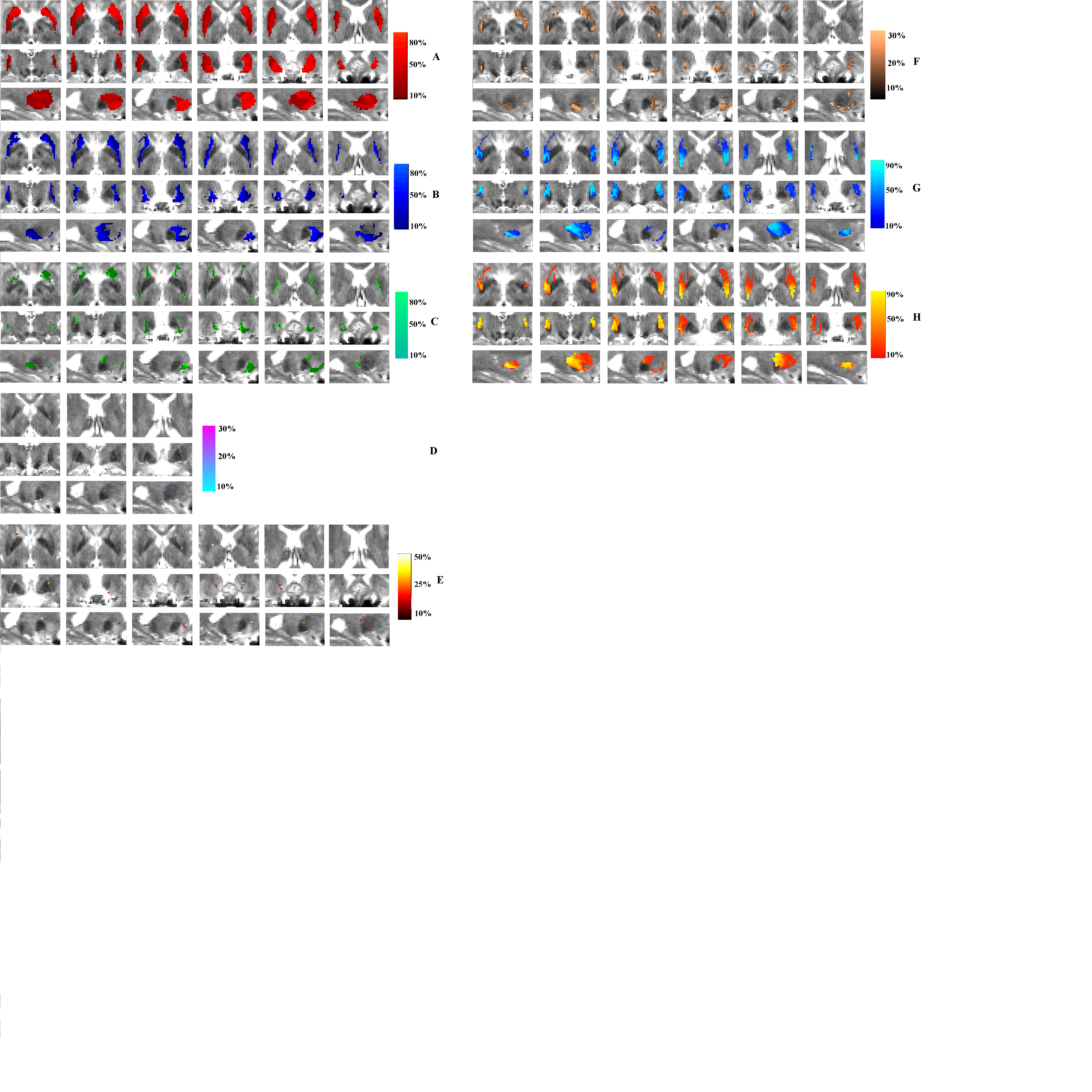

### Supplement 3 (GPe Connectivity Map)

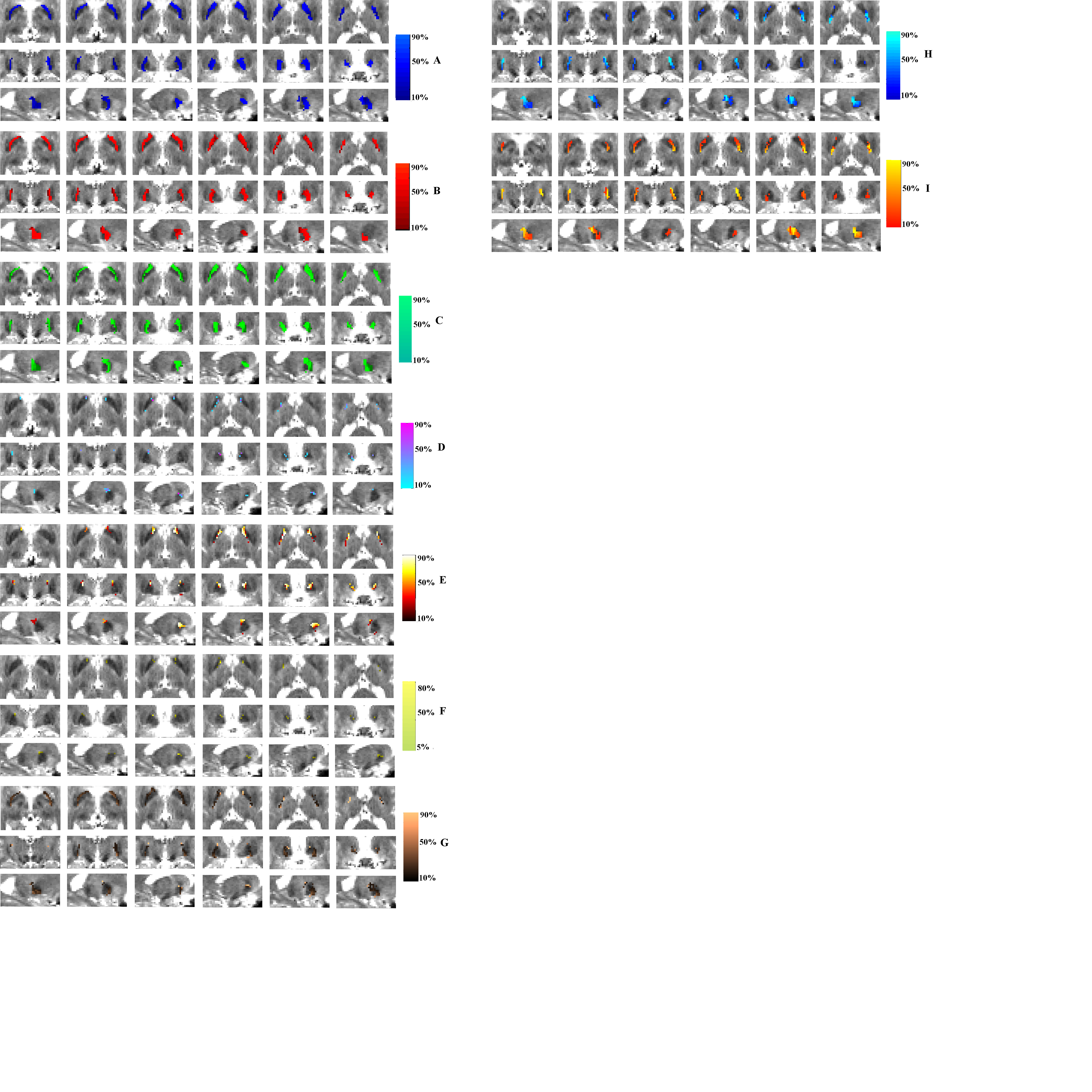

### Supplement 4 (GPi Connectivity Map)

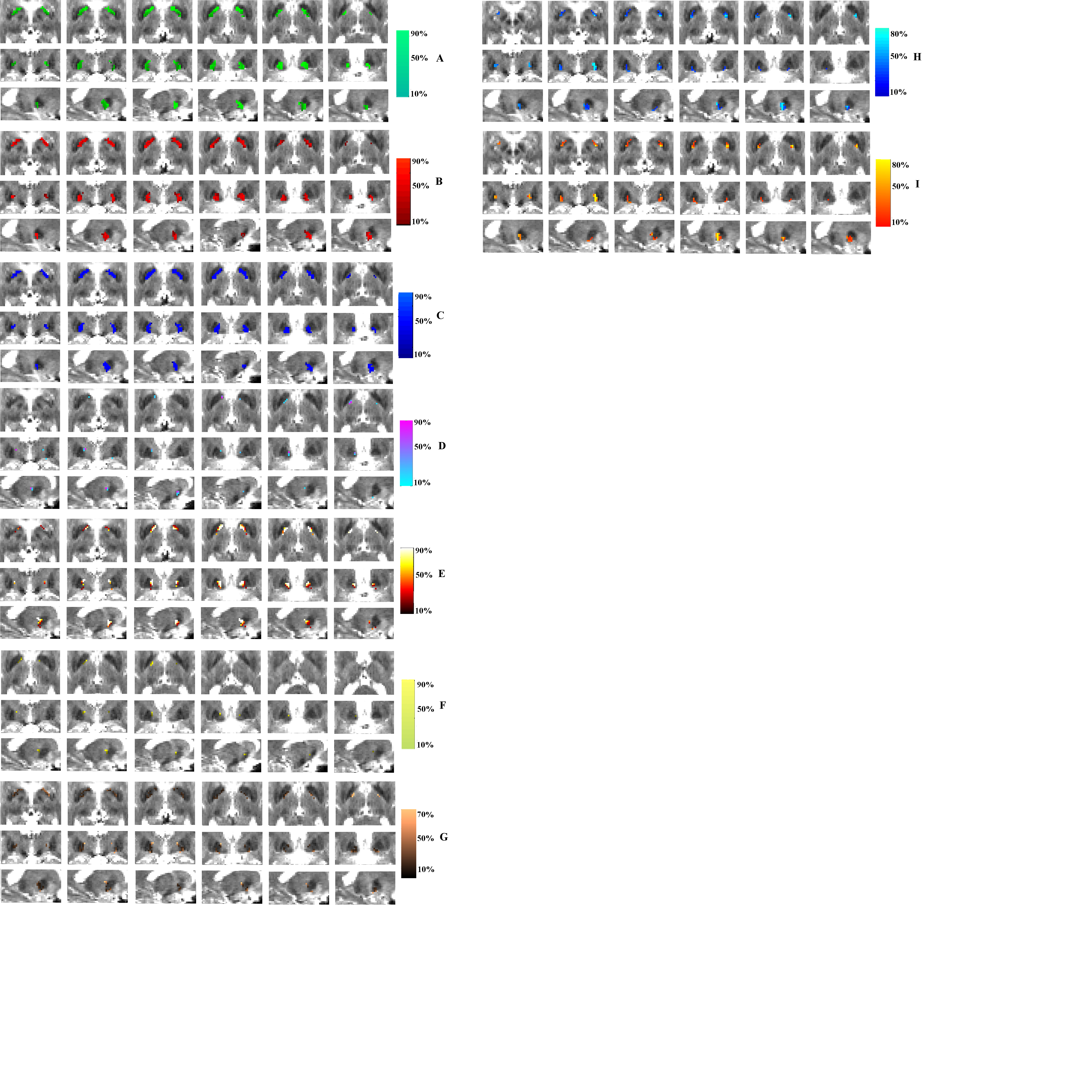

### Supplement 5 (SNc Connectivity Map)

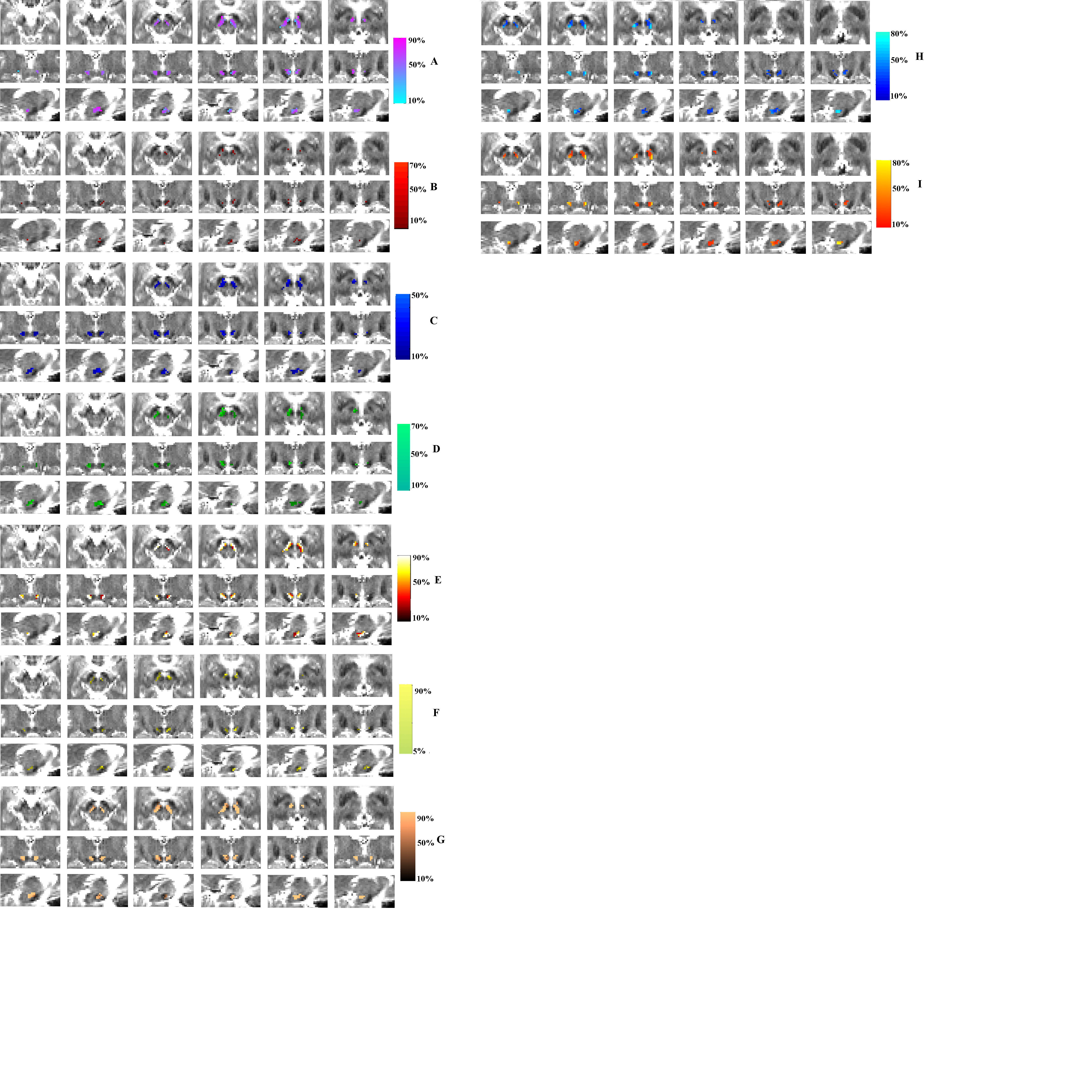

### Supplement 6 (SNr Connectivity Map)

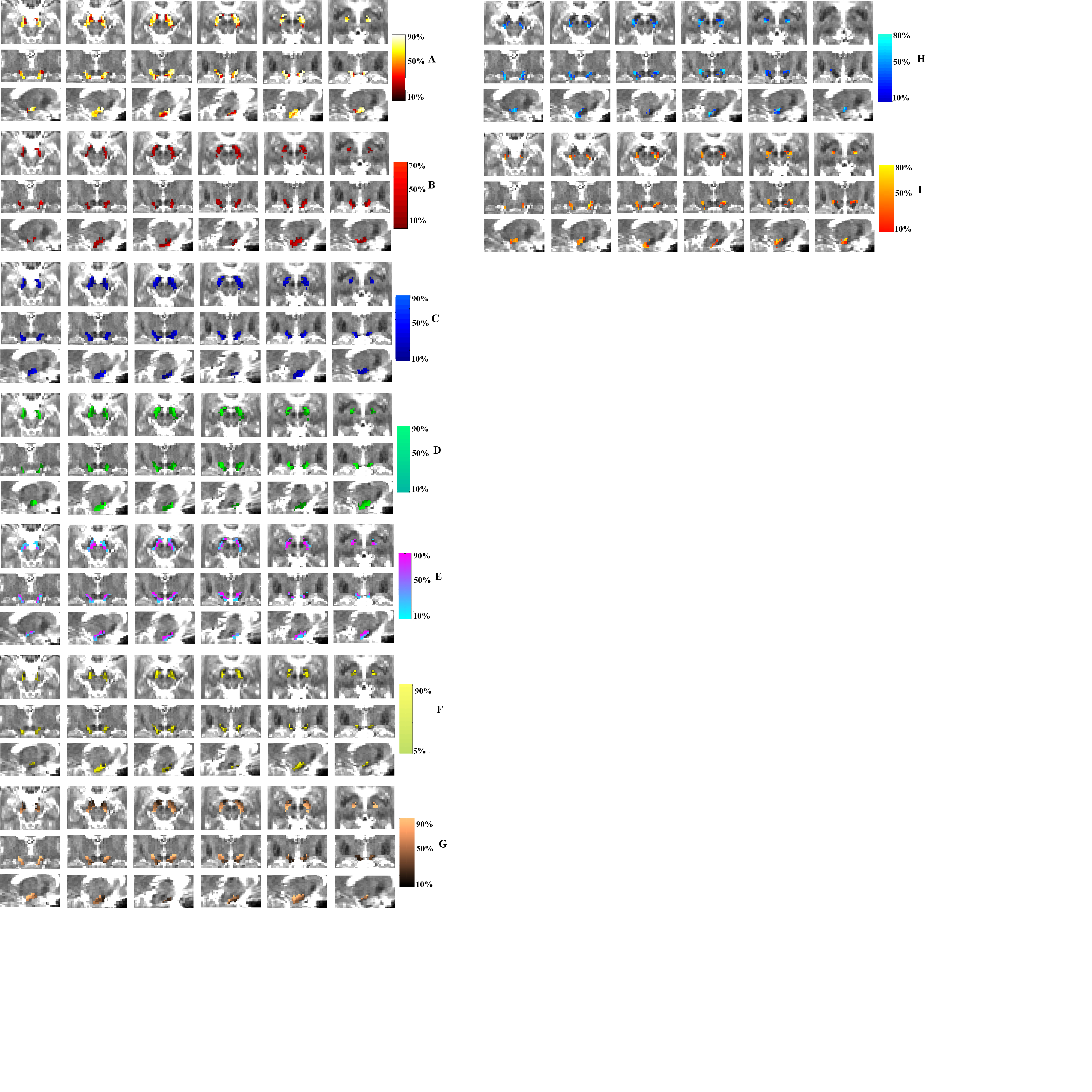

### Supplement 7 (SNc Connectivity Map)

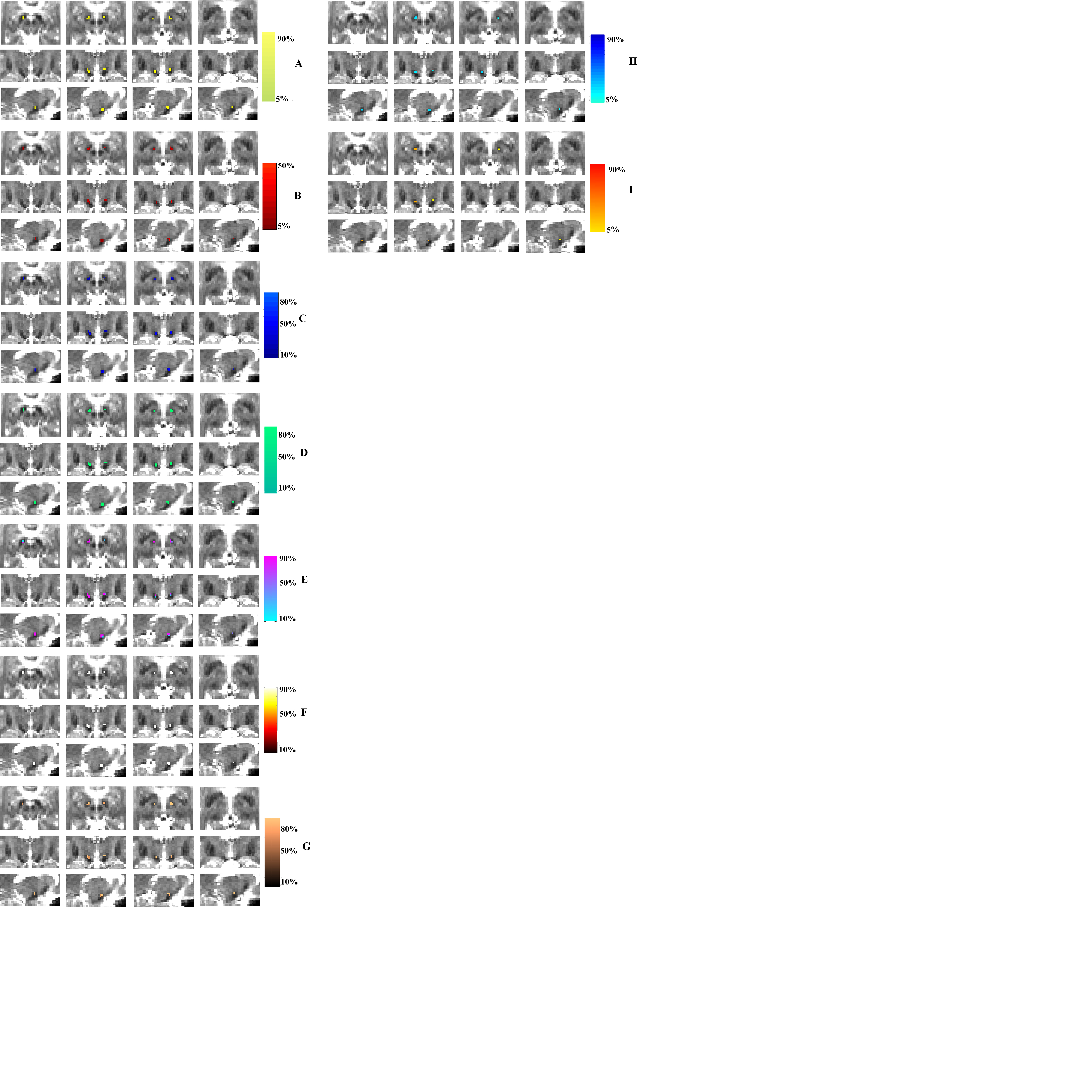

### Supplement 8 (Tha Connectivity Map)

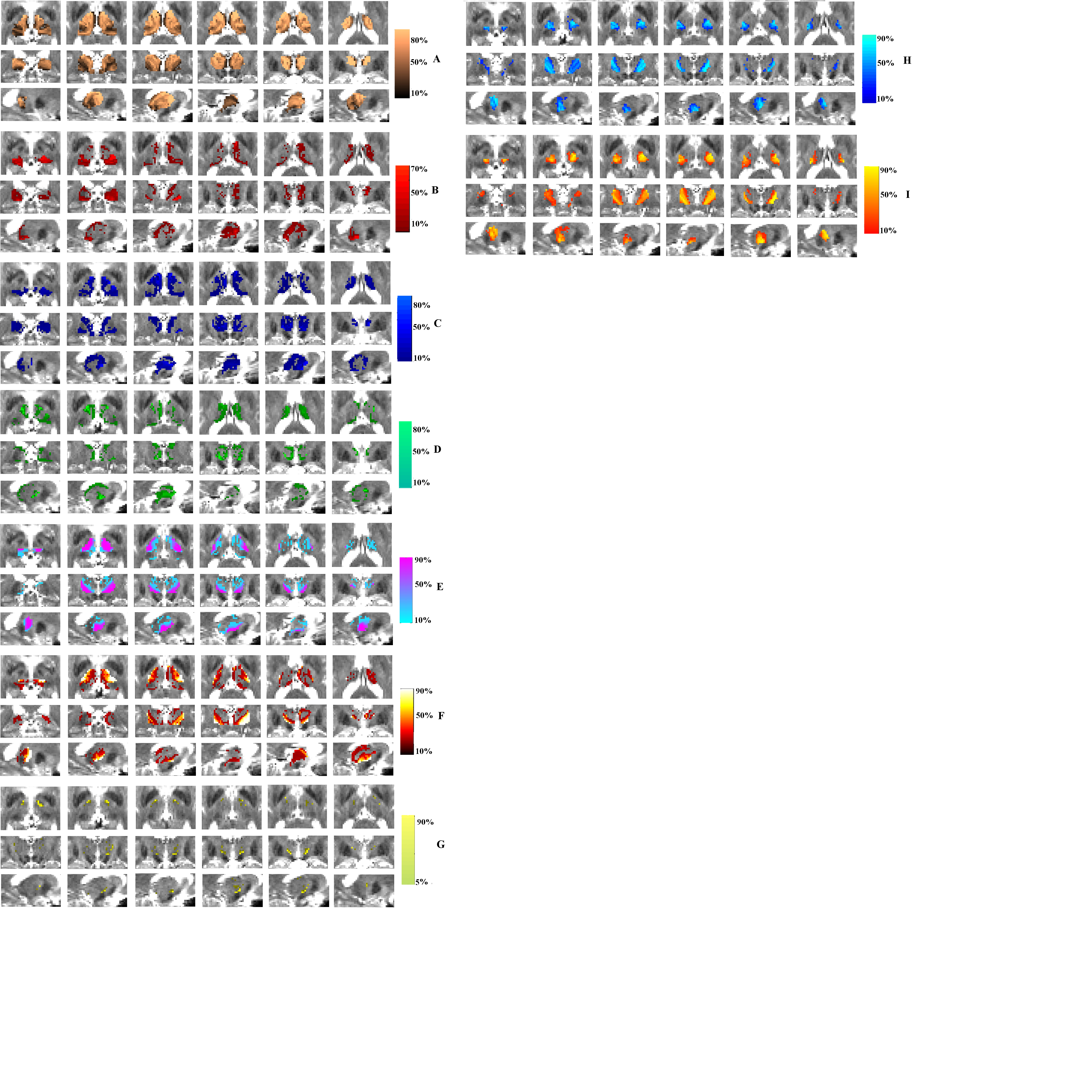
